# Altered conformational sampling along an evolutionary trajectory changes the catalytic activity of an enzyme

**DOI:** 10.1101/2020.02.03.932491

**Authors:** Joe A. Kaczmarski, Mithun C. Mahawaththa, Akiva Feintuch, Ben E. Clifton, Luke A. Adams, Daniella Goldfarb, Gottfried Otting, Colin J. Jackson

## Abstract

Several enzymes are known to have evolved from non-catalytic proteins such as solute-binding proteins (SBPs). Although attention has been focused on how a binding site can evolve to become catalytic, an equally important question is: how do the structural dynamics of a binding protein change as it becomes an efficient enzyme? Here we performed a variety of experiments, including double electron-electron resonance (DEER), on reconstructed evolutionary intermediates to determine how the conformational sampling of a protein changes along an evolutionary trajectory linking an arginine SBP to a cyclohexadienyl dehydratase (CDT). We observed that primitive dehydratases predominantly populate catalytically unproductive conformations that are vestiges of their ancestral SBP function. Non-productive conformational states are frozen out of the conformational landscape *via* remote mutations, eventually leading to extant CDT that exclusively samples catalytically relevant compact states. These results show that remote mutations can reshape the global conformational landscape of an enzyme as a mechanism for increasing catalytic activity.

## Introduction

Solute-binding proteins (SBPs) comprise a large superfamily of extra-cytoplasmic receptors that are predominantly involved in sensing and the uptake of nutrients, including amino acids, carbohydrates, vitamins, metals and osmolytes^1–4^. The SBP fold consists of two α/β domains linked by a flexible hinge region that mediates conformational change in solution, with hinge-bending and hinge-twisting motions moving the two domains together (closed state) and apart (open state; **Figure 1a**). Ligands bind at the cleft between the two domains and stabilize the closed state by forming bridging interactions between the two domains and the hinge region^1^. The ligand-induced open-to-closed conformational switch is important for the function of SBPs in ATP-binding cassette transporter systems^1,5,6^, tripartite ATP-independent periplasmic transporter sytems^7^ and signalling cascades^4^.

**Figure 1.**
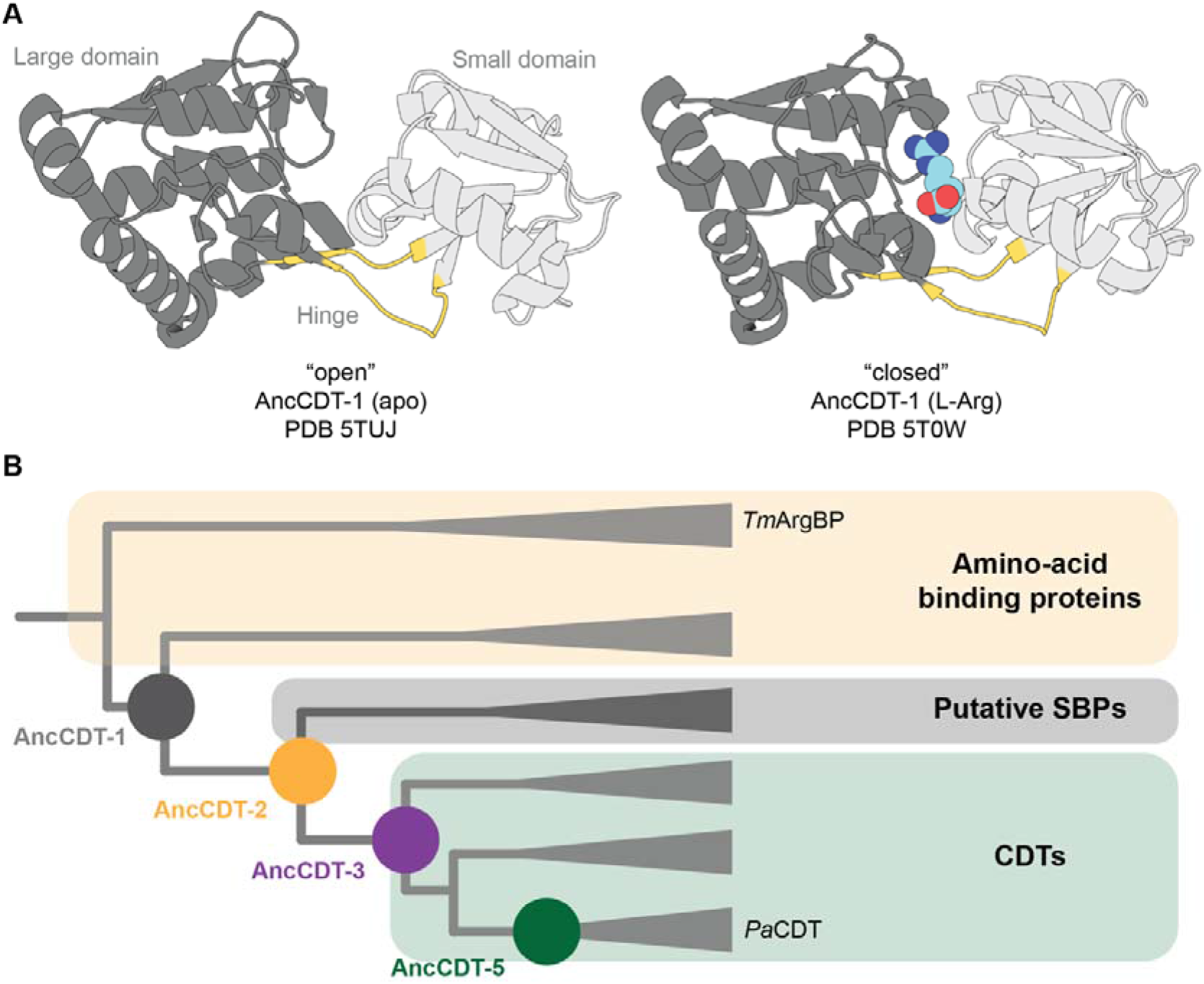
The conformational change of SBPs and the evolution of *Pa*CDT. (A) X-ray crystal structures of SBPs that are specialized for binding solutes, such as AncCDT-1 (shown), typically capture open ligand-free (left, PDB 5TUJ) and closed liganded (right, PDB 5T0W) states. (B) Schematic drawing (not to scale) of the phylogenetic tree used for ancestral sequence reconstruction in Clifton et al., 2018^40^, which highlights the evolutionary relationship between the polar amino acid binding proteins (e.g. *Thermotoga maritima* L-arginine binding protein, *Tm* ArgBP), AncCDT-1, AncCDT-3, AncCDT-5 and *Pa*CDT. Clades are collapsed. Figure adapted from Clifton et al., 2018^40^.

Although the open conformation is the ground state for most ligand-free SBPs^1,5,8–19^, numerous SBPs also sample semi-closed and closed states in the absence of ligands. For example, X-ray crystallography, nuclear magnetic resonance (NMR), molecular dynamics (MD) simulation, double electron-electron resonance (DEER, a.k.a. PELDOR) and Förster resonance energy transfer (FRET) studies on the maltose-binding protein (MBP)^20–22^, glucose-galactose-binding protein^23,24^, histidine-binding protein^25^, ferri-bacillibactin-binding protein^26^, glutamine binding protein^27,28^ and choline/acetylcholine binding protein^29^ demonstrate sampling of both open and closed conformations in the absence of ligands. The intrinsic open/closed equilibrium is fundamental to determining the binding affinity^21,23,30,31^ and binding promiscuity^5,32^ of SBPs, and controlling the transport activity of SBP-associated systems^5,28^. Indeed, the extent of the open/closed motion differs between SBPs^1^, and the function of an SBP can be changed by mutations that alter conformational sampling, without changing the architecture of the SBP-ligand interface^19,21,30^.

While most SBPs are non-catalytic binding proteins, a small fraction of proteins within the periplasmic binding protein-fold superfamily have evolved enzymatic activity^33–37^. One example is cyclohexadienyl dehydratase (CDT), which is closely related to the polar and cationic amino acid binding proteins, such as the arginine binding protein (ArgBP)^37,38^. Like ArgBP, CDT adopts a periplasmic type-II SBP fold, but instead of binding amino acids it catalyzes the Grob-like fragmentation of prephenate and L-arogenate to form phenylpyruvate and L-phenylalanine, respectively^39^. We previously used ancestral sequence reconstruction to show that the trimeric CDT from *Pseudomonas aeruginosa* (*Pa*CDT) plausibly evolved from a monomeric cationic amino acid binding protein ancestor (AncCDT-1) *via* a series of intermediates (AncCDT-2 to AncCDT-5)^40^ (**Figure 1b**). While AncCDT-1 and AncCDT-2 did not display any enzymatic activity and appeared to be binding proteins, low-level CDT activity became detectable in two alternative reconstructions of AncCDT-3(P188/L188) (*k*_cat_/*K*_M_ ~10^1^/10^2^ s^−1^ M^−1^) and increased along the trajectory towards the efficient extant enzyme *Pa*CDT (*k*_cat_/*K*_M_ ~10^6^ s^−1^ M^−1^). Despite the large difference in catalytic efficiency, AncCDT-3 and *Pa*CDT share 14/15 inner-shell residues, including all catalytic residues, but only about 50 % amino acid sequence identity over the rest of the protein (the outer shells). This suggests that these outer-shell substitutions must be substantially responsible for the ~10^5^-fold increase in catalytic specificity. For example, the remote P188L substitution in AncCDT-3 was shown to result in a 27-fold increase in *k*_cat_/*K*_M_^40^.

In this study, we investigated two plausible explanations for how remote mutations could have led to an increase in catalytic activity along this evolutionary trajectory: they may have (i) altered the sampling of rotamers of active site residues, such as the general acid Glu173, thereby changing the structure and character of the active site and controlling the configuration of active site residues, and/or (ii) altered the equilibrium between open and closed states of the protein to minimize sampling of the catalytically unproductive open state. To test these hypotheses, we used a combination of protein crystallography, MD simulations and DEER distance measurements on protein variants into which we incorporated the unnatural amino acid *p*-azidophenylalanine (AzF) to allow biorthogonal conjugation, making it specific even in the presence of native cysteine residues, to a propargyl-DO3A-Gd(III) tag^41^. The Gd(III)–Gd(III) DEER measurements were carried out at W-band frequency (94.9 GHz), affording superior sensitivity and measurements free from orientation selection and multi-spin effects^42,43^. The results from these experiments showed that the ancestral proteins, which all displayed either no catalytic activity (AncCDT-1) or very high *K*_M_ values that were outside the physiologically relevant substrate concentration (AncCDT-3, AncCDT-5), significantly or even predominantly sampled open states, including a wide-open state that is not observed in the extant and efficient enzyme *Pa*CDT. Finally, structural analysis revealed that multiple changes to intra- and intermolecular (*via* oligomerization) interaction networks have shifted the conformational equilibrium towards more compact states along the evolutionary trajectory.

## Materials and Methods

### Protein numbering convention

Residues of all proteins are referred to using the equivalent position in the reconstructed ancestral sequences (i.e. not including N-terminal tags), as described in previous work^40^ (see sequence alignment, **Figure S1**). As a consequence, residue numbers mentioned in this work differ from the residue numbers in the published PDB files of 3KBR (shift of −25), 5HPQ (−11), 6BQE (−11), 5T0W (−11), 5TUJ (−11), 5JOS (−11) and 6OKI (−11).

### Materials

The genes encoding AncCDT-1, AncCDT-3/P188, AncCDT-3/L188, AncCDT-5 and *Pa*CDT (UniProt: Q01269, residues 26–268) in pDOTS7 vectors (a derivative of pQE-28L, Qiagen) were obtained from previous work^40^. The Δ*pheA* strain of *E. coli* K-12 from the Keio collection (strain JW2580-1) was obtained from the Coli Genetic Stock Center (Yale University, CT).

### Prephenate dehydratase assay

For prephenate dehydratase assays, protein expression and purification and spectrophotometric assays were performed as described previously^40^. Proteins were expressed in Δ*pheA* cells to exclude the possibility of contamination with endogenous prephenate dehydratase. Proteins were purified by immobilized metal affinity chromatography (IMAC) and size-exclusion chromatography (SEC), and prephenate dehydratase activity was measured by end-point enzyme assays monitoring formation of phenylpyruvate by absorbance at 320 nm.

### Measurement of oligomeric state

IMAC-purified, His_6_-tagged AncCDT-1, AncCDT-3/P188, AncCDT-5 and *Pa*CDT (expressed in Δ*pheA* cells) were transferred into size-exclusion chromatography-multi angle light scattering (SEC-MALS) buffer (20 mM Na_2_HPO_4_, 150 mM NaCl, pH 7.4) and concentrated using an Amicon centrifuge filter (10 kDa molecular weight cut-off, MWCO). Samples (100 μL) of each protein were loaded at 10 mg/mL onto a pre-equilibrated Superdex 200 10/300 GL size-exclusion column (GE Healthcare) attached to multi-angle light scattering (DAWN HELEOS 8; Wyatt Technologies) and refractive index detection (Optilab rEX; Wyatt Technologies) units. A flow rate of 0.5 mL/min was used. The multi-angle detectors were normalized using monomeric bovine serum albumin (Sigma, A1900). A *dn*/*dc* value of 0.186 g^−1^ was used for each sample. The data were processed using ASTRA (Wyatt Technologies). Data were collected from a single experiment (n = 1).

### Protein sample preparation for DEER experiments

For DEER experiments, genes encoding each protein were cloned into pETMCSIII^44^ and expressed with an N-terminal His_6_-tag followed by a TEV cleavage site. In order to minimize tag side-chain dynamics, the crystal structures of the native proteins were inspected in PyMOL (The PyMOL Molecular Graphics System, Version 2.0 Schrödinger, LLC.) to select mutation sites where the sidechains of AzF residues were predicted to populate single χ_1_-angle conformations. Amber stop codons were introduced at these positions (**Table S1**) by a modified QuikChange protocol using mutant T4 DNA polymerase^45^.

All proteins for DEER experiments were expressed in RF1-free *E. coli* strain B-95.ΔA cells^46^ co-transformed with a plasmid for the aminoacyl-tRNA synthetase/tRNA pair^47^ for incorporation of AzF (Chem-Impex, USA). To minimize the amount of AzF required (provided at 1 mM), 1 L of cell-culture grown in LB medium was concentrated to 300 mL (by centrifugation and resuspension) before induction with 1 mM IPTG. Expression was conducted at 37 °C and limited to 3 hours after IPTG induction to minimize the chemical reduction of AzF. Cells were harvested by centrifugation at 5,000 × *g* for 15 min and lysed by passing two times through a French Press (SLM Aminco, USA) at 830 bars. The lysate was centrifuged at 13,000 × *g* for 30 min and the filtered supernatant was loaded onto a 5 mL Ni-NTA HisTrap HP column (GE Healthcare, USA) equilibrated with binding buffer (50 mM Tris-HCl, pH 7.5, 150 mM NaCl, 5 % w/v glycerol). The protein was eluted using elution buffer (50 mM Tris-HCl, pH 7.5, 150 mM NaCl, 5 % w/v glycerol, 300 mM imidazole) and fractions were analyzed by 12 % SDS-PAGE.

To liberate and remove ligand molecules that may have bound during protein expression, the preparation of a second set of protein samples included a denaturation and on-column refolding step. After cell lysis, guanidinium hydrochloride was added to the supernatant to a final concentration of 6 M. The filtered solution was then loaded onto a Ni-NTA column and washed with binding buffer containing 8 M urea to remove any bound ligand. Refolding was achieved using a gradient of binding buffer with decreasing amounts of urea overnight at a flow rate of 0.5 mL/min. Refolded protein was eluted as described above.

His_6_-tags were removed by digestion with His_6_-tagged TEV protease^48^ in 1:100 molar ratio overnight at 4 °C in TEV cleavage buffer (50 mM Tris-HCl, pH 8.0, 300 mM NaCl and 1 mM β-mercaptoethanol). Finally, the cleaved His_6_-tag and TEV protease were removed by passing through the Ni-NTA column (pre-equilibrated with binding buffer) prior to ligation with the propargyl-DO3A tag loaded with Gd(III).

Click reactions with the propargyl-DO3A-Gd(III) tag were performed overnight at room temperature as described previously (**Figure S2a**)^49^ and samples were exchanged into EPR buffer (20 mM Tris-HCl, pH 7.5 in D_2_O) using a Amicon Ultra-15 filter unit (10 kDa MWCO). The tagging yields were assessed by mass spectrometry, using an Elite Hybrid Ion Trap-Orbitrap mass spectrometer coupled with an UltiMate S4 3000 UHPLC (Thermo Scientific, USA).

DEER measurements were done on samples containing 100 μM protein and 20 % w/v glycerol-d_8_ in D_2_O. A 1.5-fold molar excess of L-arginine was added to one sample of refolded AncCDT-1 prior to the DEER measurement.

### DEER measurements

DEER measurements were conducted at 10 K on a home-built pulse EPR spectrometer operating at W-band (94.9 GHz)^50,51^. For the doubly labelled AncCDT-1 samples, a variant of the standard four-pulse DEER sequence (π/2(ν_obs_) – τ_1_– π(ν_obs_) – (τ_1_ + t) – π(Δν_pump_) – (τ_2_ −t) – π(v_obs_) – τ_2_ – echo) was used^52,53^. The DEER echo was observed at 94.9 GHz with π/2 and π pulses of 15 ns and 30 ns, respectively, and the field was positioned at the peak of the Gd(III) spectrum. The pump pulse was replaced by two consecutive chirp pulses produced by an arbitrary waveform generator^52–54^ which were positioned on both sides of the centre of the Gd(III) spectrum with frequency ranges of 94.5-94.8 GHz and 95-95.3 GHz, respectively. The length of each chirp pulse was 96 ns. The cavity was tuned at 94.9 GHz with a maximal microwave amplitude, ω_1_/2π, of approximately 30 MHz. Other parameters used were τ_1_ = 350 ns, a repetition rate of 0.8 ms, and τ_2_ typically ranging between 4 and 7 μs for samples with shorter and longer distances, respectively. The delay t was varied during the experiments. For the rest of the samples, a reverse DEER (rDEER) sequence (π/2(ν_obs_) – τ_1_ – π(ν_obs_) – (τ_1_ − t) – π(ν_pump_) – (τ_2 +_ t) – π(ν_obs_) – τ_2_ – echo) was used to reduce artefacts^52^. The chirp pump pulses and the positioning of the observe and the pump pulses were unchanged. In this implementation, both τ_2_ and τ_1_ typically ranged between 4 and 7 μs for shorter and longer distances, respectively.

The data were analyzed using DeerAnalysis^55^ and distance distributions were obtained using Tikhonov regularization. The regularization parameter was chosen either by the L-curve criterion or by visual inspection in order to obtain reasonable fits. Estimation of uncertainties in distance distributions due to background correction were obtained using the validation option in DeerAnalysis^55^.

### Crystallization of AncCDT-5 and structure determination

A solution of TEV-cleaved AncCDT-5 (expressed in BL21(DE3) cells and purified by IMAC as described above) was further purified by SEC using a HiLoad 16/600 Superdex 75 pg column (GE Healthcare, USA) in 50 mM Tris-HCl, pH 7.5, 20 mM NaCl, and was concentrated to 30 mg/mL using an Amicon Ultra-15 filter unit (10 kDa MWCO). The protein was crystallized using the vapor diffusion method at 18 °C. Crystallization screens were set up using the PACT premier HT-96 C10 screen from Molecular Dimensions (Newmarket, USA) with drops containing 200 nL reservoir solution and 200 nL AncCDT-5 solution. AncCDT-5 crystals formed in 28 days in 20 % w/v PEG 6000, 0.2 M MgCl_2_, 0.1 M HEPES, pH 7.0.

Crystals were cryoprotected (30 % w/v PEG 6000, 0.2 M MgCl_2_, 0.1 M HEPES pH 7.0) and frozen in liquid nitrogen. Data were collected at 100 K on the MX2 beamline of the Australian Synchrotron. The data were indexed and integrated in XDS^56^ and scaled in AIMLESS^57^. The structure was solved by molecular replacement in Phaser^58^ using the two domains of *Pa*CDT as search models (PDB 3KBR small domain = residues 122-223; large domain = residues 27-121 and 224-258). The structure was refined by real space refinement in Coot^59^ and using REFMAC^60^ and phenix.refine^61^. Data collection and refinement statistics are given in **Table S2**. Residues in the AncCDT-5 structure were numbered according to the equivalent positions in AncCDT-1 (PDB 5TUJ)^40^ and the coordinates deposited in the Worldwide Protein Data Bank (PDB 6OKI).

### Molecular dynamics (MD) simulations

500 ns simulations were performed in Desmond^62^ (in Schrödinger 2019−11) using the OPLS3e force field^63^. These simulations were initiated from the *Pa*CDT–acetate trimer (PDB 5HPQ, residues 13-250 in PDB, n=1), unliganded AncCDT-1 (PDB 5T0W, chain A, residues 14-246, n=3), AncCDT-3/L188 (PDB 5JOS, residues 12-247, n=3), AncCDT-3/P188 (n=3) and AncCDT-5 (PDB 6OKI, residues 13-245, n=3) structures, with all small molecules removed. The AncCDT-3/P188 structure was obtained by making the L188P mutation and local minimization in Maestro (Schrödinger Release 2019-1: Maestro, Schrödinger, LLC, New York, NY, 2019). Desmond was used to add N-terminal acetyl caps and C-terminal amide caps to each structure and for energy minimization of the protein structures. Each protein was solvated in an orthorhombic box (15 Å buffer periodic boundary) with SPC water molecules and the systems were neutralized using Na^+^ or Cl^−^ ions. Energy minimization was achieved using a hybrid method of the steepest descent algorithm and the limited-memory Broyden–Fletcher– Goldfarb–Shanno algorithm (maximum of 2,000 iterations and a convergence threshold of 1 kcal/mol/Å). The system was relaxed using the default relaxation procedure in Desmond. For production MD simulations of the NPT ensemble, the temperature was maintained at 300 K using a Nosé–Hoover thermostat (relaxation time = 1.0 ps) and the pressure was maintained at 1.01 bar (relaxation time = 2.0 ps) using a Martyna–Tobias– Klein barostat. Otherwise, default Desmond options were used. Following relaxation of the system, each simulation was run for 500 ns. Distances between the α-carbons of the tagged residues and the radii of gyration were calculated using the ProDy package on snapshots sampled every 0.1 ns^64^. In addition, representative snapshots (with a range of Cα–Cα distances) corresponding to the closed, open and wide-open states (if sampled) were selected for modelling of Gd(III)–Gd(III) distances. Allocation of states (closed/open/wide-open) was done based on Cα–Cα measurements. One snapshot, which was initially allocated as open based on its Cα–Cα distance, could also be grouped as wide-open based on principal component analysis (PCA, performed in ProDy), and was therefore included in both groups.

### Modelling of propargyl-DO3A-Gd(III) tags and Gd(III)–Gd(III) distances

To calculate predicted Gd(III)–Gd(III) distances of different conformations, propargyl-DO3A-Gd(III) tags were modelled onto the crystal structures of AncCDT-1 (PDB 5T0W, 5TUJ), AncCDT-3/L188 (PDB 5JOS), AncCDT5 (PDB 6OKI), *Pa*CDT (PDB 3KBR, 5HPQ) and selected MD snapshots. To do this, the mutation tool in PyMOL was used to identify the side-chain dihedral angles χ_1_ and χ_2_ of a phenylalanine residue that produced the least amount of steric clashes with the rest of the protein; these angles were used when modelling the tag. The angles χ_9_ and χ_10_ (**Figure S2b**) were fixed to −140° and 70°, respectively, to allow coordination of the metal ion by the nearest nitrogen of the triazole ring. The angle χ_6_ was set to 180°. As in previous work with a closely related cyclen tag^49^, setting χ_6_ to 180° yielded a better correlation between experimental and back-calculated Gd(III)–Gd(III) distances compared with setting χ_6_ to 0° (data not shown).

## Results

### The evolution of prephenate dehydratase activity

We previously showed that while AncCDT-1 binds L-arginine with comparable affinity to extant L-arginine binding proteins, the P188 AncCDT-3 reconstruction (hereafter AncCDT-3/P188), AncCDT-5 and *Pa*CDT have all lost the ancestral ability to bind proteogenic amino acids^40^. Instead, along with the AncCDT-3/L188 AncCDT-3 reconstruction, they can rescue the growth of a phenylalanine auxotroph of *E. coli* (Δ*pheA*) grown in the absence of L-phenylalanine, indicating their capacity to deliver the dehydratase activity required to synthesize L-phenylalanine from L-arogenate, or the precursor phenylpyruvate from prephenate^40^. Furthermore, these variants exhibited prephenate dehydratase activity *in vitro*, which increased about 10^5^-fold (*k*_cat_/*K*_M_) between AncCDT-3/P188 and *Pa*CDT. In this work, we have additionally assessed the dehydratase activity of AncCDT-5, which proved to be intermediate between the AncCDT-3 variants and the extant enzyme *Pa*CDT (**Table 1**, **Figure S3**). Although the *k*_cat_ value of AncCDT-5 (4 s^−1^) exceeds the level observed in the AncCDT-3 variants (~10^−2^ s^−1^) almost as much as *Pa*CDT (18 s^−1^), the *K*_M_ of AncCDT-5 remains ~15-fold higher than that of *Pa*CDT (277 μM *vs.* 19 μM). This value is well above the likely physiological concentration of the substrate (the intracellular concentrations of molecules in this biosynthetic pathway are about 14–18 μM in Gram-negative bacteria^65^).

**Table 1.** Kinetic parameters for prephenate dehydratase activity of CDT variants related to this study.*

| Protein | $K_{\text{M}}$ ($\mu\text{M}$) | $k_{\text{cat}}$ (s<sup>-1</sup>) | $k_{\text{cat}}/K_{\text{M}}$ (M<sup>-1</sup> s<sup>-1</sup>) |
| --- | --- | --- | --- |
| AncCDT-1** | ND | ND | ND |
| AncCDT-3/P188** | 1830 ± 190 | (1.04 ± 0.07) $\times 10^{-2}$ | 5.67 ± 0.70 |
| AncCDT-3/L188** | 294 ± 27 | (4.58 ± 0.30) $\times 10^{-2}$ | 155 ± 18 |
| AncCDT-5 | 277 ± 4 | 4.1 ± 0.2 | (1.5 ± 0.1) $\times 10^4$ |
| *Pa*CDT** | 18.7 ± 2.9 | 18.4 ± 0.7 | (9.8 ± 1.6) $\times 10^5$ |

ND, not detected.
\* Results are reported as mean ± standard error for $K_{\text{M}}$ and $k_{\text{cat}}$ . Errors for $k_{\text{cat}}/K_{\text{M}}$ were obtained by propagation.
\*\*<sup>40</sup>

### Remote mutations affect active site configurations and catalytic efficiency

Sequence and structural analysis of AncCDT-3/L188 and *Pa*CDT revealed that although their catalytic specificity differ by ~10^5^ M^−1^ s^−1^, they share 14 of 15 residues in the active/substrate binding site (the only difference being a conservative Thr80Ser substitution)^40^. Comparison between the structures of AncCDT-3/L188 and *Pa*CDT revealed that the catalytic general acid Glu173 adopts a rotameric state in AncCDT-3/L188 that differs from that observed in *Pa*CDT (**Figure 2a**)^40^. This conformational difference appears to be caused by the neighbouring Tyr177Gln mutation that occurs on the branch between AncCDT-3 and AncCDT-5; the presence of Tyr177 in AncCDT-3 prevents Glu173 from adopting the catalytically competent conformation observed in *Pa*CDT^40^. Additionally, this mutation causes coupled conformational changes in the neighbouring residues Phe156 and Met167, which leads to further changes in the shape and electrostatic character of the active site. These findings are analogous to those from recent work on enzyme variants produced through directed evolution and computational design that show how second-shell mutations affect enzyme activity by constraining the conformational sampling of catalytic residues^66,67^.

**Figure 2.**
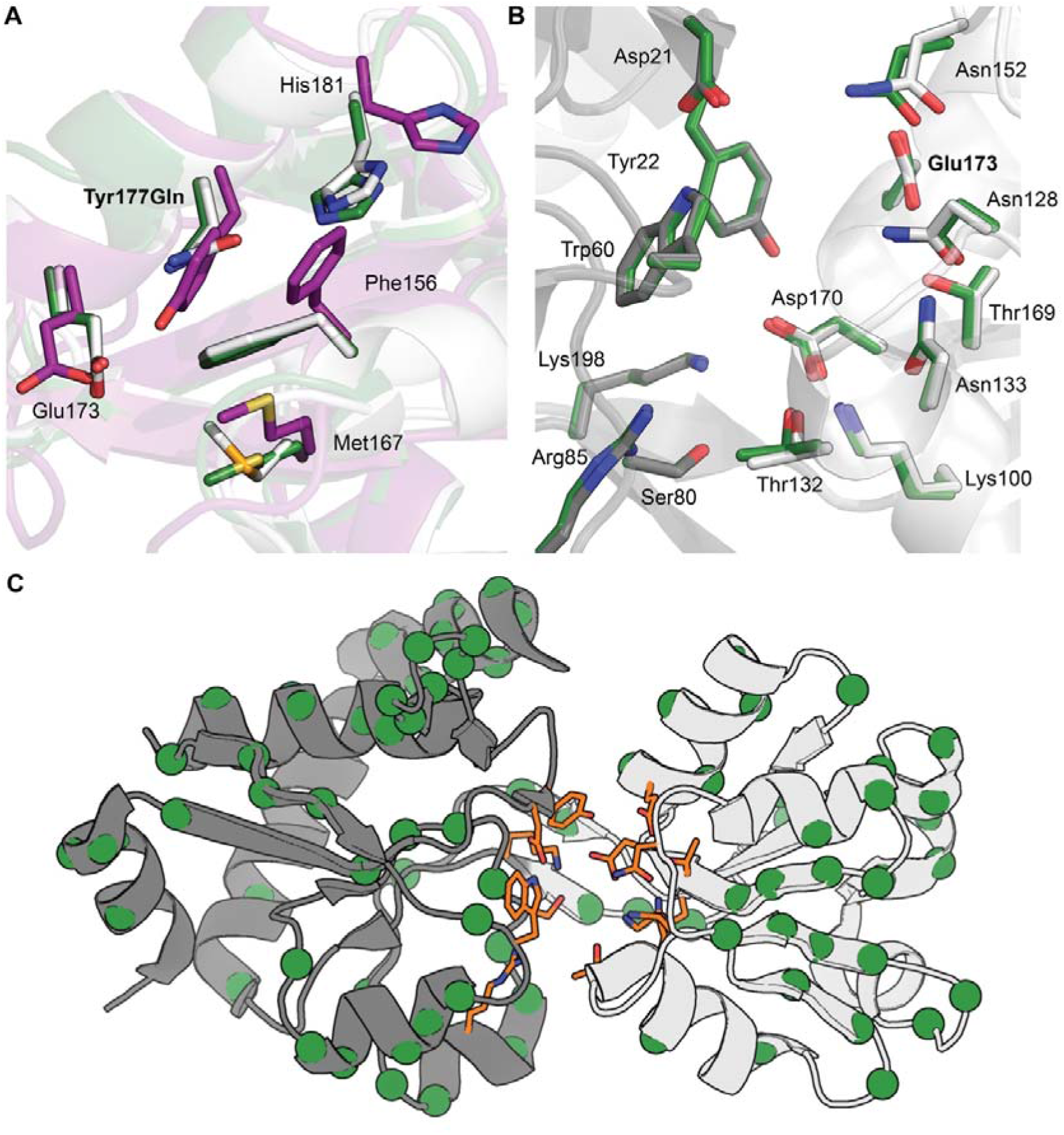
Crystal structures of AncCDT-3/P188, AncCDT-5 and *Pa*CDT. (A) Structural overlay of the small domains of AncCDT-3/L188 (purple, PDB 5JOS), AncCDT-5 (green) and *Pa*CDT (white, PDB 3KBR). The Tyr177Gln substitution that occurs between AncCDT-3 and AncCDT-5 contributes to the repositioning of Glu173. Structural alignment of the active sites of *Pa*CDT (PDB 3KBR, white/grey) and AncCDT-5 (green). The small domain (white, residues 99–194) and large domain (grey, residues 1–98 and 195–236) of AncCDT-5 were individually aligned with the corresponding domains of *Pa*CDT. The side chains of the 15 inner-shell residues identified by Clifton et al., 2018^40^ are shown as sticks (Gly131 not shown), highlighting identical active site residues and side-chain conformations between AncCDT-5 and *Pa*CDT. The HEPES molecules present in the crystal structures are omitted for clarity. The locations of the substitutions (green spheres) between AncCDT-5 and *Pa*CDT are shown projected onto the structure of *Pa*CDT (PDB 3KBR). Inner-shell residues are shown as orange sticks.

To better understand how the mutations along this evolutionary trajectory affect activity, we also solved a 1.37 Å X-ray crystal structure of AncCDT-5 (**Table S2**), which shares 15/15 residues in the active/substrate binding site with *Pa*CDT. In contrast to the comparison between AncCDT-3 and *Pa*CDT, a structural alignment of AncCDT-5 and *Pa*CDT (PDB 3KBR) reveals that these two proteins not only share identical binding sites in terms of amino acid composition, but that these residues also adopt identical conformations (**Figure 2b**). Since these active/binding sites are identical, the ~15-fold decrease in *K*_M_ and ~5-fold increase in *k*_cat_ of the dehydratase activity must be attributed to amino acid substitutions that are outside of the binding site. The sequences of AncCDT-5 and *Pa*CDT differ at 98 positions. Excluding the 9 amino acid C-terminal extension of *Pa*CDT, the remaining 89 positions are evenly distributed around the SBP fold (**Figure 2c**), making it difficult to rationalize how individual mutations contribute to changes in catalytic profiles. Instead, it is possible that some of the substitutions may be affecting catalysis by altering the open/closed dynamics of these proteins.

### Solution dynamics of the amino acid binding protein, evolutionary intermediates and *Pa*CDT

Next, we investigated whether the accumulation of substitutions between AncCDT-1 and *Pa*CDT at positions remote from the active site could have affected catalytic activity by altering the open/closed dynamics of these proteins. To do so, we used DEER distance measurements and MD simulations to investigate whether the distribution of open and closed solution states differed between the ancestral amino acid binding protein (AncCDT-1), the intermediate low-activity CDTs (AncCDT-3, AncCDT-5) and the efficient extant enzyme, *Pa*CDT.

#### AncCDT-1

First, we confirmed that AncCDT-1 was monomeric (>95%) in solution using size-exclusion chromatography-multi angle light scattering (SEC-MALS; (**Figure S4a**). We then assessed the solution conformations of AncCDT-1 using DEER distance measurements with Gd(III)-tags introduced through unnatural amino acid (UAA) mutagenesis. Amber stop codons were used to incorporate the UAA AzF into the small (Lys138) and large (Gln68) domains of AncCDT-1. These sites were chosen because they (i) are known to be located in regions of the protein that undergo substantial changes in their relative distance upon ligand binding (**Figure S5**)^40^, (ii) are located on rigid α-helices, (iii) are surrounded by residues that will minimize unwanted tag motions *via* packing and (iv) are solvent exposed, which minimizes the risk that mutagenesis and tagging at these positions will significantly affect the protein conformations. The mutant protein heterologously expressed in soluble form from *E. coli*, yielding ~15 mg of purified protein per litre of cell culture. To remove any co-purified ligands, on-column protein denaturation and refolding was performed to obtain ligand-free samples. This is referred to as refolded AncCDT-1. The AzF residues were modified with propargyl-DO3A-Gd(III) tags as previously described (**Figure S2a**)^41,49^ and mass spectrometry analysis indicated tag ligation yields of >80 % at each site.

DEER measurements of refolded AncCDT-1-propargyl-DO3A-Gd(III) revealed a distance distribution with two dominant peaks with maxima corresponding to Gd(III)– Gd(III) distances of 3.2 nm and 4.4 nm (**Figure 3a**, **Table S3, Figure S5a,b**). When saturating concentrations of L-arginine were added to the refolded sample, the distance distribution revealed an almost complete shift to the shorter distance (**Figure 3a**), as expected for a ligand-induced open-to-closed transition. Together, these data indicate that the Gd(III)–Gd(III) distance varies by ~1.2 nm between the closed state and any open conformations and that the DEER approach employed here can be used to differentiate between distinct conformational substates of a SBP. Experiments with natively purified protein rather than unfolded/refolded protein showed similar results (**Figure S5a,b**), except that a greater proportion of the protein adopted the closed state (3.2 nm Gd(III)–Gd(III) distance) in the absence of exogenous L-arginine than in the sample subjected to unfolding and renaturation. This was expected, as the protein is known to co-purify from *E. coli* with some ligand bound, which stabilizes the closed conformation^40^.

**Figure 3.**
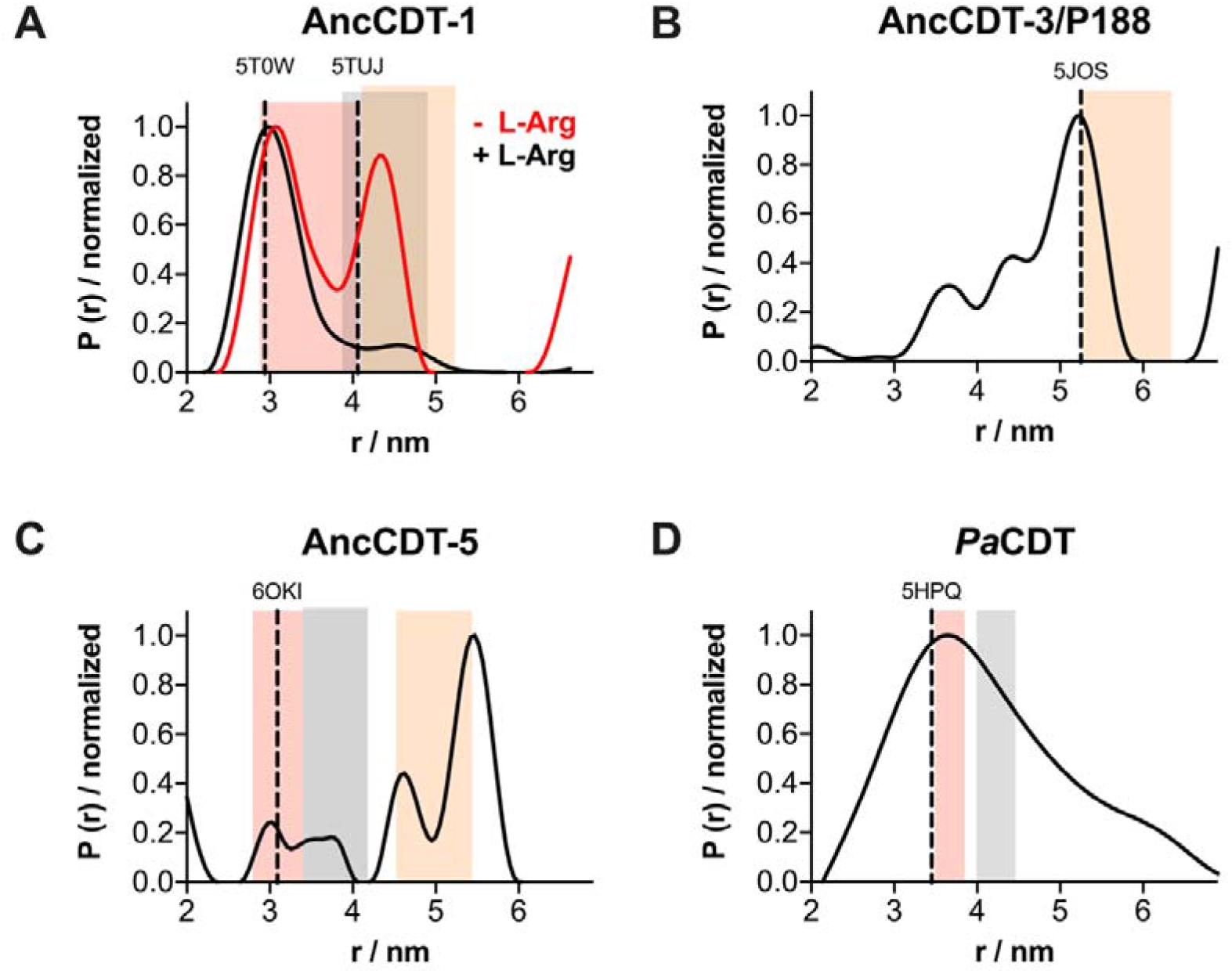
Solution dynamics of AncCDT-1, AncCDT-3/P188, AncCDT-5 and *Pa*CDT probed by DEER. Gd(III)–Gd(III) distance distributions for samples of propargyl-DO3A-Gd(III)-tagged (A) refolded AncCDT-1 (−/+ L-Arg, tagged at positions 68 and 138), (B) natively purified AncCDT-3/P188 (tagged at positions 68 and 138), (C) natively purified AncCDT-5 (tagged at positions 68 and 138) and (D) natively purified *Pa*CDT (tagged at positions 68 and 139 and diluted by a factor of 10 using unlabelled *Pa*CDT). Distance distributions were obtained using DeerAnalysis^55^. The original DEER data and DEER form factor traces are shown in Figures S5-7. Vertical dashed lines represent the Gd(III)–Gd(III) distances estimated by modelling propargyl-DO3A-Gd(III) tags onto the crystal structures of each protein. The shaded areas represent the range of calculated Gd(III)–Gd(III) distances when propargyl-DO3A-Gd(III) tags are modelled onto a number of individual MD snapshots representing the closed (red), open (grey) and wide-open (orange) states.

We also performed experiments on AncCDT-1 that had been tagged at two sites on either the large domain (positions 68 and 219, **Figure S5c**) or small domain (positions 138 and 161; **Figure S5d**). These each showed a peak in the distance distribution that were in agreement with Gd(III)-Gd(III) distance predictions based on crystal structures of AnCDT-1 (**Figure S5g,h**). These results confirm that the two distances measured for refolded AncCDT-1 are due to rigid body motions of the two domains, rather than any intra-domain flexibility or tag dynamics.

We next sought to correlate the distance changes observed through DEER to structural changes in the protein. We previously solved the crystal structure of AncCDT-1 bound to L-arginine (PDB 5T0W), which has a Cα–Cα distance between Gln68 and Lys138 of 2.6 nm (equivalent to a Gd(III)–Gd(III) distance of ~2.9 nm, **Figure S5e**). We also solved a structure of the protein in an open state (PDB 5TUJ) in which the Cα–Cα distance between Gln68 and Lys138 is 3.4 nm (Gd(III)–Gd(III) distance of ~4.1 nm, **Figure S5f**). This crystal structure presents a snapshot of an open conformation, but additional open states may be accessible to the protein in solution. To probe these, we also explored the solution dynamics of AncCDT-1 using MD simulations initiated from the closed X-ray crystal structure of AncCDT-1 (PDB 5T0W) with the ligand, L-arginine, removed. These were conducted solely to identify conformational substates that can be accessed on the ns-μs timescale (i.e. easily accessible, low-energy states), not to obtain quantitative predictions of the solution dynamics or the relative occupancy of each state. The simulations suggest that AncCDT-1 can easily access three dominant conformational substates (**Figure 4a**, **Table S3**): the closed state corresponding to PDB 5T0W (68-138 Cα–Cα distance range of 2.5-2.9 nm; equivalent to a Gd(III)–Gd(III) distance range of ~2.9-3.4 nm), an intermediate open state corresponding to PDB 5TUJ (68-138 Cα–Cα distance 3.2-3.7 nm; equivalent to a Gd(III)–Gd(III) distance range of ~3.9-4.9 nm) and an additional wide-open state (68-138 Cα–Cα distance 3.6-4.2 nm; equivalent to a Gd(III)–Gd(III) distance range of ~4.1-5.2 nm).

**Figure 4.**
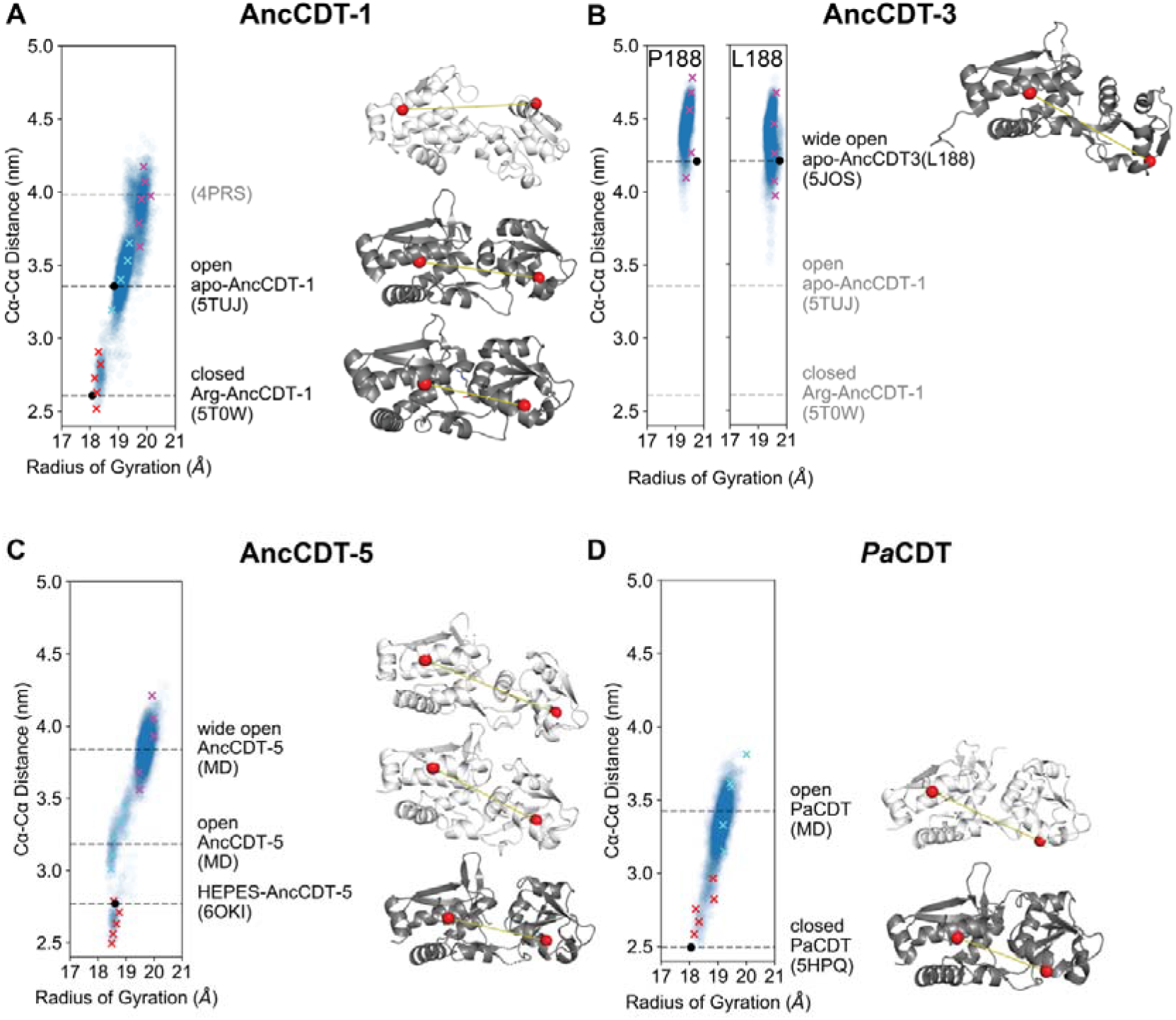
Conformational substates captured by MD and crystallography. Conformational substates of (A) AncCDT-1, (B) AncCDT-3, (C) AncCDT-5 and (D) *Pa*CDT captured by MD and X-ray crystallography. The plot of Cα**–**Cα distances between tagged residues *vs.* radius of gyration obtained from MD simulation snapshots highlights populations (or lack thereof) of closed, open and wide-open conformational states. Each data point represents a single frame of the MD simulation (sampled every 0.1 ns). Crosses represent snapshots of closed (red), open (cyan) and wide-open (magenta) states that were used to model Gd(III)–Gd(III) distances. Representative structures of each of these states are shown on the right and highlight the extent of the open/closed transition, with distances between Cαs (red spheres) of tagged sites shown by yellow lines. Dark grey structures are crystal structures (corresponding to black dots on plot), while white structures are MD snapshots.

The results obtained by DEER suggest that the dominant states are indeed the closed state and an open conformation closely related to the open state observed through protein crystallography, but which might also include the wide-open state, as there is significant overlap between the modelled Gd(III)–Gd(III) distances in the snapshots obtained from MD. Although we do not have crystallographic evidence that AncCDT-1 samples the wide-open state, analogous wide-open states have been observed in crystal structures of extant L-arginine binding proteins, such as that from *Thermotoga maritima* (*Tm*ArgBP, PDB 4PRS)^38^, which has a Cα–Cα distance of 4.0 nm between positions equivalent to 68-138 in AncCDT-1 (which is similar to the 3.6-4.2 nm range observed for the wide-open state in AncCDT-1 MD simulations). Indeed, it has been suggested that the specific open states of SBPs observed crystallographically can be dictated by crystal packing interactions^8–11^. The wide-open state thus is a part of the conformational landscape for L-arginine binding proteins, although we could not unambiguously detect it in our DEER experiments on AncCDT-1 owing to overlap with the open state. Altogether, the data from DEER, protein crystallography and MD demonstrate that AncCDT-1 can sample both open and closed states in the absence of ligands, and that ligand binding shifts the conformational equilibrium towards the closed state.

#### AncCDT-3 and AncCDT-5

Considering the significant difference in dehydratase activity between AncCDT-3, AncCDT-5 and *Pa*CDT, as well as our inability to fully rationalize these differences through protein crystallography, we next used DEER and MD to investigate the solution conformational distributions of these proteins. Like AncCDT-1, SEC-MALS indicated that AncCDT-3/P188 and AncCDT-5 were predominantly monomeric in solution (>95%), as reported previously^40^ (**Figure S4b,c**). We tagged the small and large domains of AncCDT-3/P188 and AncCDT-5 at sites 68 (large domain) and 138 (small domain) in the same way as AncCDT-1 and performed DEER experiments (**Figure 3b,c**, **Figure S6a,b**). For natively purified AncCDT-3/P188, the DEER distance distributions showed a clear maximum that corresponded to a Gd(III)–Gd(III) distance of ~5.2 nm (**Figure 3b**, **Figure S6a**). Likewise, AncCDT-5 displayed the most prominent distance-distribution peak at ~5.4 nm (**Figure 3c**, **Figure S6b**). Similar results were obtained for the refolded proteins (**Figure S6a,b**), where the observed shifts are within experimental error.

We previously solved a crystal structure of AncCDT-3/L188 and found a conformation analogous to the wide-open conformation observed in the MD simulation of AncCDT-1 (PDB 5JOS)^40^. In this structure, the Q68-R138 Cα–Cα distance is 4.2 nm (Gd(III)–Gd(III) distance: ~5.3 nm). In triplicate 500 ns MD simulations initiated from the open AncCDT-3/L188 and AncCDT-3/P188 structures (after modelling the L188P mutation), the protein did not sample the closed state at all across the combined 1.5 ms of simulation time and the wide-open state was the major conformation (**Figure 4b**), exhibiting a Cα–Cα distance range of 4.0-4.8 nm (Gd(III)–Gd(III) distance range: 5.0-6.3 nm).

Unlike the wide-open crystal structure of AncCDT-3/L188, the 1.38 Å crystal structure of AncCDT-5 determined in the present work adopts the closed conformation (a HEPES molecule is bound in the active site), with a Q68-R138 Cα–Cα distance of 2.8 nm (Gd(III)–Gd(III) distance: 3.1 nm). Beginning from this closed state, we again performed triplicate 500 ns MD simulations after removing the HEPES molecule. The simulations predict that the protein rapidly adopts a wide-open conformation, with a Q68-R138 Cα–Cα distance range of 3.6-4.2 nm (Gd(III)–Gd(III) distance range: 4.5-5.4 nm) that is analogous to the wide-open conformation seen in the MD simulations of AncCDT-1, the crystal structure of the L-arginine binding protein from *T. maritima,* and the crystal structure of AncCDT-3 (**Figure 4c**). An intermediate state, analogous to the open crystal structure of unliganded AncCDT-1 was also briefly sampled, which corresponded to a Q68-R138 Cα–Cα distance range of 3.0–3.4 nm (Gd(III)–Gd(III) distance range: 3.4–4.2 nm). The combination of the DEER, crystallographic and MD data for the experiments performed on AncCDT-3 and AncCDT-5 suggest that both proteins predominantly access a wide-open state in solution. Smaller peaks in the distance distribution could correspond to minor fractions of intermediate states but are too small to be unambiguous.

#### Pa*CDT*

We next aimed to probe the open/closed dynamics of *Pa*CDT. SEC-MALS experiments confirmed that *Pa*CDT is primarily homotrimeric in solution^40^, with a small population (< 5 %) of higher order or aggregated species (**Figure S4d**), which complicates the measurement of intra-monomer distances by DEER experiments. To confirm the existence of the tagged trimer, we first measured samples with propargyl-DO3A-Gd(III) tags at single AzF residues of each *Pa*CDT monomer, at either the large domain (position 68) or the small domain (position 139). The maxima of the DEER distance distributions measured were consistent with those expected for the trimeric structure observed by X-ray crystallography (3.5 nm *vs.* 3.3 nm for proteins tagged at position 68, 5.4 nm *vs*. 4.6 nm for samples tagged at position 139; **Figure S7a,b**). This indicated that the trimeric structure was preserved under the conditions of the DEER experiments. When we attempted to measure the open/closed distribution *via* labelling at sites on both the large and small domain (68 and 139, respectively) the observed Gd(III)–Gd(III) distance distribution was very broad as was expected due to multiple inter-chain Gd(III)–Gd(III) distances in a trimer (**Figure S7c**). In order to extract the inter-domain distance from a single *Pa*CDT monomer, a protein sample labelled at sites 68 and 139 was diluted 10-fold with unlabelled protein and allowed to equilibrate, with the aim of obtaining a mixture in which there would be a ~10 %, 1 % and 0.1 % chance of encountering a trimer consisting of 1, 2 or 3 tagged monomers, respectively. DEER experiments on samples prepared in this way yielded a significantly more narrow peak that indicated 3.5 nm was the predominant Gd(III)–Gd(III) distance (**Figure 3d, Figure S7d**).

We again used protein crystallography and MD simulation to identify stable conformations of *Pa*CDT. There are three crystal structures, one with HEPES bound at the active site (Structural Genomics Project, PDB 3KBR) and two with acetate bound at the active site^40^. The highest resolution acetate-bound structure (PDB 5HPQ) and the HEPES-bound structure (PDB 3KBR) display Q68-A139 Cα–Cα distances of 2.5 nm (Gd(III)–Gd(III) distance: ~3.5 nm). A MD simulation initiated from the trimeric X-ray crystal structure of acetate-*Pa*CDT (PDB 5HPQ), with acetate removed, indicated that the monomers can sample both closed (Cα–Cα distance range of 2.5–3.0 nm; equivalent to a Gd(III)–Gd(III) distance range of ~3.5–3.9 nm) and open (Cα– Cα distance range of 3.2–3.8 nm; equivalent to a Gd(III)–Gd(III) distance range of ~4.0–4.5 nm) conformations, but there was no evidence of a wide-open conformation (**Figure 4d**). The Gd(III)–Gd(III) distances predicted for the closed and open states notably fall either side of the broad peak centred at 3.9 nm. This could be consistent with a broad distribution of rapidly interconverting open and closed states that have been snap-frozen, in contrast the AncCDT-1 DEER analysis, which displayed two distinct peaks corresponding to similar distances.

All three crystal structures of *Pa*CDT display closed conformations, which leave limited access to the active site. To allow exchange of substrate and product, the enzyme must also populate open conformations, which are likely related to those obtained in the MD simulations. Interestingly, however, the wide-open conformation, which appears in MD simulations of AncCDT-1, AncCDT-3 and AncCDT-5, was not significantly sampled. This is, again, consistent with the DEER measurements, where the prominent peaks at ~5.4 nm in AncCDT-3 and AncCDT-5 were absent for *Pa*CDT. Altogether, the combined use of DEER, crystallography and MD simulations suggests that the wide-open conformation observed in the ancestral reconstructions of CDT (computationally observed in all variants and empirically observed in AncCDT-3, AncCDT-5 and orthologs of AncCDT-1) has been largely eliminated from the conformational landscape of the extant *Pa*CDT. This provides a molecular explanation for the observed change in *K*_M_ between AncCDT-5 and *Pa*CDT; despite their identical substrate binding sites when closed, the open ligand-binding state of *Pa*CDT likely has higher affinity for the substrate as it is much closer to the conformation of the Michaelis complex than the wide-open state that is predominantly populated in AncCDT-3 and AncCDT-5.

### Structural basis for the increased dehydratase activity of *Pa*CDT

Having established that the conformational sampling of *Pa*CDT is different from AncCDT-5, we compared the structures of AncCDT-5 and *Pa*CDT to ascertain the molecular basis for this change. Structural comparison between the two proteins revealed three regions with substantial differences (unlike the substrate binding site) that could collectively stabilize the closed state and/or destabilize the wide-open state. First, in comparison to AncCDT-5, which terminates at Lys235 (although it is not present in clear electron density), *Pa*CDT has an additional nine amino acids at the C-terminus (RWPTAHGKL), of which the first 4 and 6 residues produce clear electron density in crystal structures of *Pa*CDT (PDB 5HPQ and 6BQE, respectively) (*30*). As shown in **Figure 5a**, these amino acids interact extensively with both the small and large domains, substantially increasing the number of van der Waals and hydrogen bond interactions between the two domains in the closed and open, but not wide-open, conformation.

**Figure 5.**
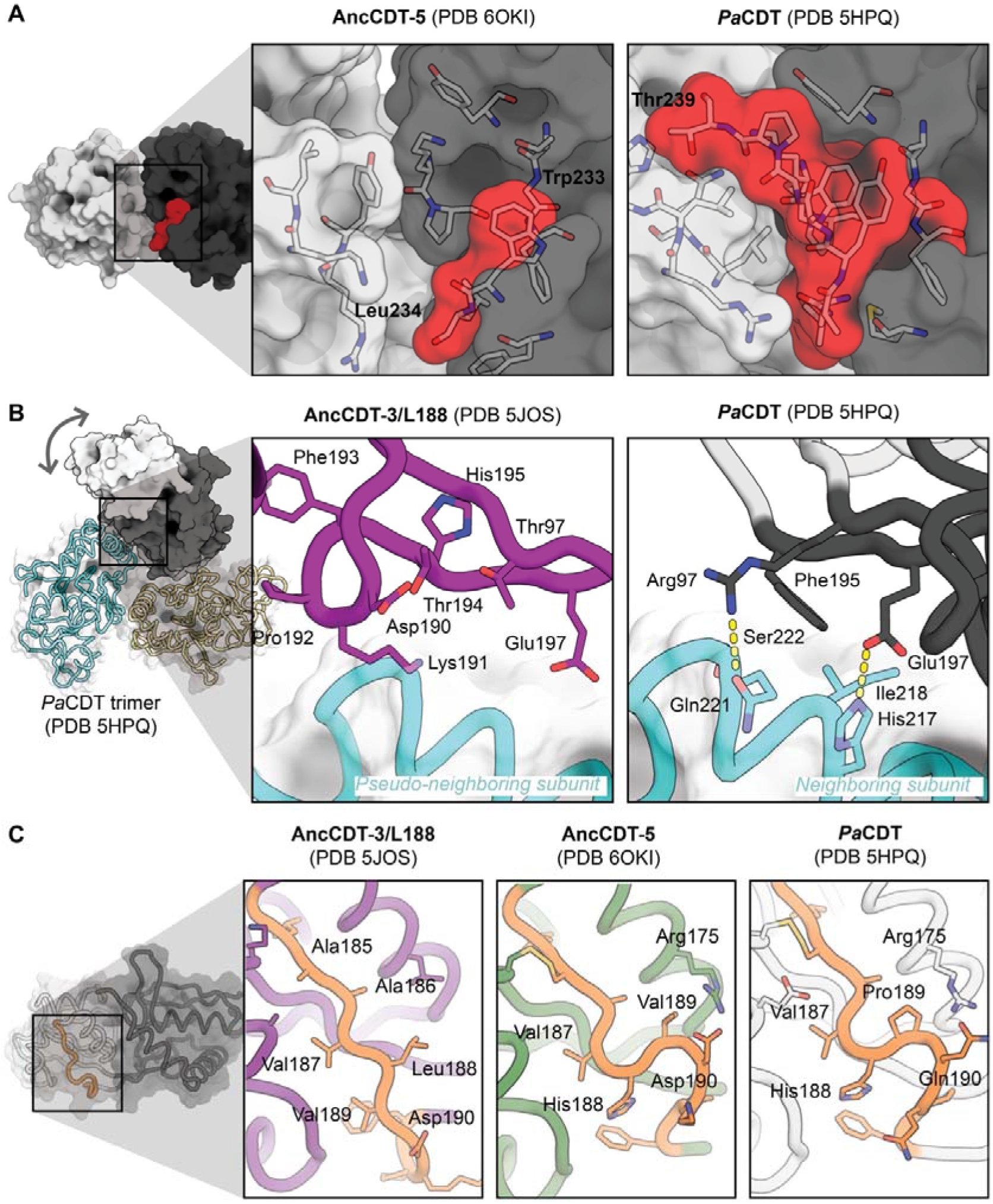
Structural basis for change in conformational sampling. (A) Comparison of the crystal structures of AncCDT-5 (left) and *Pa*CDT (5HPQ, right) highlight how the C-terminal extension (residues beyond 232 are highlighted in red) in *Pa*CDT facilitates additional interactions between the small (white) and large (grey) domains. (B) *Pa*CDT is a trimer in solution, and analysis of the crystal structures of *Pa*CDT (e.g. 5HPQ, shown here) reveals several interactions between the hinge region of chain A with the large domain of chain B that are likely to prevent the wide-open conformation being sampled. These include packing of Phe195 against the neighbouring subunit, and polar interactions between neighbouring residues. Overlay of the crystal structure of AncCDT-3/L188 (purple, PDB 5JOS) further highlights how the wide-open state is likely to be incompatible with the trimeric arrangement of *Pa*CDT. (C) The flexibility of the hinge region is known to control the open/closed dynamics of SBPs. Sequence and structural comparison of these regions in AncCDT-3/L188 (purple, PDB 5JOS), AncCDT-5 (green, PDB 6OKI) and *Pa*CDT (white, PDB 5HPQ) highlight that this is an intrinsically flexible region, which is likely affected by mutations.

Second, one of the most substantial structural differences between AncCDT-5 and *Pa*CDT is the oligomerization of *Pa*CDT, which forms a trimer (**Figure 5b**). In this trimer, the small domain remains free to move, allowing fluctuation between the closed and open states. However, there are inter-subunit contacts at the hinge region that appear to preclude some of the conformational changes expected in the wide-open state. Specifically, the crystal structure of the wide-open conformation of AncCDT-3 shows that the hinge region with His195 at the centre is fully solvent exposed and extended. The equivalent region of *Pa*CDT, however, is in a retracted conformation, with Phe195 (chain A; the equivalent position to His195 in AncCDT-3) being stabilized in a buried hydrophobic pocket formed by Ile218 (chain A) and the non-polar region of Ser222 (chain A), as well as the non-polar regions of two hydrogen-bonded pairs, Arg97 (chain A):Gln221 (chain B) and Glu197 (chain A):His217 (chain B). The positioning of these residues and interactions with the neighbouring subunit would likely stabilize *Pa*CDT in the closed and open, while limiting the sampling of wide-open states.

Finally, for the proteins to adopt the wide-open state, the region between Val187 and Pro191 must undergo a conformational change, from the tightly kinked conformation seen in the closed structures of AncCDT-5 and *Pa*CDT (PDB 5HPQ, 6BQE; it is in an altered conformation due to crystal packing interactions in 3KBR), to an extended conformation seen in the crystal structure of the wide-open conformation of AncCDT-3 (**Figure 5c**). In AncCDT-3/5, the region between Val187 and Pro191 is relatively flexible, consisting of polar and relatively small hydrophobic residues (Leu/Val/Glu in AncCDT-3; His/Val/Glu in AncCDT-5). In contrast, in *Pa*CDT, the central residue in the kink is mutated to proline, and the neighbouring glutamine forms a hydrogen bond to Arg186, stabilizing the kinked structure. Further support for the hypothesis that this region is important in conformational sampling and dehydratase activity can be found in the observation that (i) the P188L mutation in AncCDT-3 causes a large decrease in *K*_M_^40^, (ii) it was observed to be a mutational hotspot during the directed evolution of ancestral CDTs for increased dehydratase activity^40^ and (iii) this region has been recognized to be important for dynamics and affinity in other SBPs such as MBP^21,30^.

The combination of these three structural effects is likely sufficient to substantially change the relative conformational free energy of the closed, open and wide-open states, consistent with the DEER and MD results. Interestingly, none of the structural changes observed between AncCDT-3 and *Pa*CDT preclude *Pa*CDT adopting the open conformation, but essentially prevent it adopting the wide-open conformation. Given the large number of changes between AncCDT-5 and *Pa*CDT, it is impossible to ascribe the difference in populations of open and wide-open conformations to a single mutation or structural rearrangement. Indeed, recent work has highlighted the gradual nature by which protein conformational sampling can change, and the overlapping and often redundant effects of mutations^66^.

## Discussion

SBPs have become a model system for understanding the role of rigid-body motions in determining ligand and substrate binding affinity, specificity and kinetics. Early crystallographic studies suggested that SBPs act as simple binary switches, adopting a closed conformation when in complex with ligands and an open conformation in the ligand-free state^4^. More recently, however, NMR, FRET and MD studies have probed the intrinsic dynamics of SBPs, showing that they can adopt closed and semi-closed states in the absence of ligands^20–29^, and have provided evidence that the extent of the open-closed dynamics evolves to support specific binding properties and function. Indeed, altering the open-closed dynamics of SBPs is now also being used as an additional parameter for engineering new binding functions into SBPs^21^ and the development of novel SBP-based biosensors^68^. Such efforts, however, are complicated by the fact that very subtle changes to protein motion can alter SBP function and the structural basis for changes in these open-closed dynamics are poorly understood. Here we have expanded on this work, demonstrating that SBPs of this protein fold can access three relatively well-defined conformational substates, including a wide-open state for which the possible physiological role must still be determined. In view of the ease with which it is sampled in MD simulations and unambiguously observed by DEER experimentally in two of the native proteins we studied (AncCDT-3 and AncCDT-5), as well as the apo-state of the L-arginine binding protein from *T. maritima*^38^, it is unlikely to be an artefact. The apparently facile sampling of both closed and wide-open states could explain why FRET experiments, which have been conducted with fluorescent tags attached to either domain, sometimes report smaller changes in signal upon ligand binding than would be expected if a single open conformation was fully populated in the apo-protein^69,70^.

We previously suggested that mutations accumulated during evolution most likely increased the dehydratase activity of *Pa*CDT by shifting the conformational equilibrium away from the catalytically-unproductive open state to the catalytically-competent closed state^40^. Crystal structures and MD simulations provided some preliminary evidence that this is the case, but now we have strong quantitative evidence in solution. First, the crystal structure of AncCDT-5 revealed identical active/binding sites of AncCDT-5 and *Pa*CDT, demonstrating that amino acid substitutions remote from the active site can have very large effects on activity *via* mechanisms other than controlling the structure of the active site. Second, the dramatic shift in the open/closed equilibrium of the proteins is the main change along the evolutionary trajectory. These results show that the wide-open state, which is the lowest-energy state in AncCDT-3 and AncCDT-5 and is unlikely to play any catalytic role owing to low affinity for substrate owing to the spatial separation between the binding site/catalytic residues, appears to be depopulated in the final evolutionary step between AncCDT-5 and *Pa*CDT. Third, the extant and efficient *Pa*CDT appears to exist in a broad distribution between a closed state optimized for catalysis of the chemical step in the reaction and a moderately open state that retains the shape of the binding pocket but opens a pathway for substrate/product diffusion. Structural analysis provides a molecular explanation for this shift in conformational sampling, with a number of structural changes stabilizing the open/closed states at the expense of the wide-open state. Interestingly, this seems to have occurred at least in part through oligomerization, suggesting that evolution can favour oligomerization both for stability and activity^71^.

This work expands the scope of the DEER approach^19,72–76^ for use in this context by using Gd(III) spin labels; it is the first time that the propargyl-DO3A-Gd(III)/DEER approach has been used to study rigid-body protein dynamics and it demonstrates the excellent performance of the propargyl-DO3A-Gd(III) tag. The tag is independent of cysteine residues as it can be ligated to AzF residues and the small size, lack of net charge and hydrophilic character of the DO3A-Gd(III) complex disfavours specific interactions with the protein surface, thus increasing the reliability of computational predictions of the Gd(III)–Gd(III) distance distributions obtained with this tag^41^. This study highlights its efficiency (the DEER measurements requiring only nanogram amounts of protein) in studying the protein dynamics of a series of states along an evolutionary trajectory and how it afforded distribution widths that were sufficiently narrow to observe the co-existence of open and closed protein conformations that differ by only about 1 nm, to show the presence of two conformations simultaneously sampled in solution and to detect ligand-induced conformational changes. Importantly, the distances measured by DEER here are in good agreement with distances predicted from the crystal structures and states obtained from MD simulations.

## Conclusions

In this work, we studied the structural dynamics of a series of states along an evolutionary trajectory from an ancestral L-arginine solute binding protein (AncCDT-1) to an extant dehydratase (*Pa*CDT). The challenge of accurately defining the conformational landscape of a protein is substantial: virtually all approaches have serious limitations or are so demanding that they are not practical for studying more than a single protein. Here, by following an integrated structural biology approach involving the combined use of the three methods, the shortcomings of each approach (crystal packing forces in single crystals, imperfect force fields in MD simulations, potential overlap in the signals from different states for DEER measurements, etc.) could be compensated for. We compared crystal structures of AncCDT-3, a new structure of AncCDT-5, and *Pa*CDT, which revealed that second-shell residues alter the active site configuration between AncCDT-3 and AncCDT-5, but that AncCDT-5 and *Pa*CDT display large differences in catalytic activity despite identical active site configurations. Using a propargyl-DO3A-Gd(III) tag and W-band DEER distance measurements, we were able to experimentally assess the distribution of different conformational substates in frozen solutions along this evolutionary trajectory, with reference to structures obtained through protein crystallography and MD simulations. These data provide new insight into how proteins can evolve new functions: while the correct active site configuration must be established, this study shows that the ability of an enzyme to adopt a range of conformational states that are tailored to its catalytic role is equally important. This result contributes to the growing body of literature that suggests molecular evolution is a relatively smooth process, whereby conformational substates that are not productive are gradually depopulated while beneficial substates are enriched.

## Acknowledgements

JAK acknowledges financial support from an Australian Government Research Training Program Scholarship. BEC acknowledges financial support from a Rod Rickards PhD Scholarship. Funding by the Australian Research Council, including a Laureate Fellowship to GO, is gratefully acknowledged. DG acknowledges the support of the Minerva foundation and this research was made possible in part by the historic generosity of the Harold Perlman Family (DG). DG holds the Erich Klieger Professorial Chair in Chemical Physics.

## Accession codes

The atomic coordinates and structure factors for AncCDT-5 (PDB 6OKI) have been deposited in the Worldwide Protein Data Bank (http://wwpdb.org/).

## References

1. Davidson, A. L., Dassa, E., Orelle, C. & Chen, J. Structure, function, and evolution of bacterial ATP-binding cassette systems. Microbiol. Mol. Biol. Rev. 72, 317–364 (2008).

2. Berntsson, R. P. A., Smits, S. H. J., Schmitt, L., Slotboom, D. J. & Poolman, B. A structural classification of substrate-binding proteins. FEBS Lett. 584, 2606–2617 (2010).

3. Scheepers, G. H., Lycklama a Nijeholt, J. A. & Poolman, B. An updated structural classification of substrate-binding proteins. FEBS Lett. 590, 4393–4401 (2016).

4. Tam, R. & Saier, M. H. Structural, functional, and evolutionary relationships among extracellular solute-binding receptors of bacteria. Microbiol. Rev. 57, 320–46 (1993).

5. De Boer, M. et al. Conformational and dynamic plasticity in substrate-binding proteins underlies selective transport in ABC importers. eLife 8, 1–28 (2019).

6. Quiocho, F. A. & Ledvina, P. S. Atomic structure and specificity of bacterial periplasmic receptors for active transport and chemotaxis: Variation of common themes. Mol. Microbiol. 20, 17–25 (1996).

7. Mulligan, C. et al. The substrate-binding protein imposes directionality on an electrochemical sodium gradient-driven TRAP transporter. Proc. Natl. Acad. Sci. U. S. A. 106, 1778–1783 (2009).

8. Magnusson, U., Salopek-Sondi, B., Luck, L. A. & Mowbray, S. L. X-ray structures of the leucine-binding protein illustrate conformational changes and the basis of ligand specificity. J. Biol. Chem. 279, 8747–8752 (2004).

9. Magnusson, U. et al. Hinge-bending motion of D-allose-binding protein from *Escherichia coli*. Three open conformations. J. Biol. Chem. 277, 14077–14084 (2002).

10. Lau, A. Y. & Roux, B. The free energy landscapes governing conformational changes in a glutamate receptor ligand-binding domain. Structure 15, 1203–1214 (2007).

11. Björkman, A. J. & Mowbray, S. L. Multiple open forms of ribose-binding protein trace the path of its conformational change. J. Mol. Biol. 279, 651–664 (1998).

12. Loeffler, H. H. & Kitao, A. Collective dynamics of periplasmic glutamine binding protein upon domain closure. Biophys. J. 97, 2541–2549 (2009).

13. Chu, B. C. H., De Wolf, T. & Vogel, H. J. Role of the two structural domains from the periplasmic *Escherichia coli* histidine-binding protein HisJ. J. Biol. Chem. 288, 31409–31422 (2013).

14. Sooriyaarachchi, S., Ubhayasekera, W., Park, C. & Mowbray, S. L. Conformational changes and ligand recognition of *Escherichia coli* D-xylose binding protein revealed. J. Mol. Biol. 402, 657–668 (2010).

15. Skrynnikov, N. R. et al. Orienting domains in proteins using dipolar couplings measured by liquid-state NMR: Differences in solution and crystal forms of maltodextrin binding protein loaded with β-cyclodextrin. J. Mol. Biol. 295, 1265–1273 (2000).

16. Kang, C. H. et al. Crystal structure of the lysine-, arginine-, ornithine-binding protein (LAO) from *Salmonella typhimurium* at 2.7-Å resolution. J. Biol. Chem. 266, 23893–23899 (1991).

17. Oh, B. H. et al. Three-dimensional structures of the periplasmic lysine/arginine/ornithine-binding protein with and without a ligand. J. Biol. Chem. 268, 11348–11355 (1993).

18. Stockner, T., Vogel, H. J. & Tieleman, D. P. A salt-bridge motif involved in ligand binding and large-scale domain motions of the maltose-binding protein. Biophys. J. 89, 3362–3371 (2005).

19. Glaenzer, J., Peter, M. F., Thomas, G. H. & Hagelueken, G. PELDOR spectroscopy reveals two defined states of a sialic acid TRAP transporter SBP in solution. Biophys. J. 112, 109–120 (2017).

20. Tang, C., Schwieters, C. D. & Clore, G. M. Open-to-closed transition in apo maltose-binding protein observed by paramagnetic NMR. Nature 449, 1078–1082 (2007).

21. Seo, M. H., Park, J., Kim, E., Hohng, S. & Kim, H. S. Protein conformational dynamics dictate the binding affinity for a ligand. Nat. Commun. 5, 3724 (2014).

22. Selmke, B. et al. Open and closed form of maltose binding protein in its native and molten globule state as studied by electron paramagnetic resonance spectroscopy. Biochemistry 57, 5507–5512 (2018).

23. Ortega, G., Castaño, D., Diercks, T. & Millet, O. Carbohydrate affinity for the glucose-galactose binding protein is regulated by allosteric domain motions. J. Am. Chem. Soc. 134, 19869–19876 (2012).

24. Flocco, M. M. & Mowbray, S. L. The 1.9 Å X-ray structure of a closed unliganded form of the periplasmic glucose/galactose receptor from *Salmonella typhimurium*. J. Biol. Chem. 269, 8931–8936 (1994).

25. Chu, B. C. H., Chan, D. I., DeWolf, T., Periole, X. & Vogel, H. J. Molecular dynamics simulations reveal that apo-HisJ can sample a closed conformation. Proteins Struct. Funct. Bioinforma. 82, 386–398 (2014).

26. de Boer, M., Gouridis, G., Muthahari, Y. A. & Cordes, T. Single-molecule observation of ligand binding and conformational changes in FeuA. Biophys. J. 117, 1642–1654 (2019).

27. Feng, Y. et al. Conformational dynamics of apo-GlnBP revealed by experimental and computational analysis. Angew. Chemie - Int. Ed. 55, 13990–13994 (2016).

28. Gouridis, G. et al. Conformational dynamics in substrate-binding domains influences transport in the ABC importer GlnPQ. Nat. Struct. Mol. Biol. 22, 57–64 (2015).

29. Oswald, C. et al. Crystal structures of the choline/acetylcholine substrate-binding protein ChoX from *Sinorhizobium melilotie* in the liganded and unliganded-closed states. J. Biol. Chem. 283, 32848–32859 (2008).

30. Marvin, J. S. & Hellinga, H. W. Manipulation of ligand binding affinity by exploitation of conformational coupling. Nat. Struct. Biol. 8, 795–798 (2001).

31. Marvin, J. S. & Hellinga, H. W. Conversion of a maltose receptor into a zinc biosensor by computational design. Proc. Natl. Acad. Sci. U. S. A. 98, 4955–4960 (2001).

32. Clifton, B. E. & Jackson, C. J. Ancestral protein reconstruction yields insights into adaptive evolution of binding specificity in solute-binding protein. Cell Chem. Biol. 23, 236–245 (2016).

33. Cho, Y., Sharma, V. & Sacchettini, J. C. Crystal structure of ATP phosphoribosyltransferase from *Mycobacterium tuberculosis*. J. Biol. Chem. 278, 8333–8339 (2003).

34. Kreinbring, C. A. et al. Structure of a eukaryotic thiaminase I. Proc. Natl. Acad. Sci. U. S. A. 111, 137–142 (2014).

35. Brownlie, P. D. et al. The three-dimensional structures of mutants of porphobilinogen deaminase: toward an understanding of the structural basis of acute intermittent porphyria. Protein Sci. 3, 1644–1650 (1994).

36. Arai, R. et al. Crystal structure of MqnD (TTHA1568), a menaquinone biosynthetic enzyme from *Thermus thermophilus* HB8. J. Struct. Biol. 168, 575–581 (2009).

37. Tam, R. & Saier, M. H. A bacterial periplasmic receptor homologue with catalytic activity: cyclohexadienyl dehydratase of Pseudomonas aeruginosa is homologous to receptors specific for polar amino acids. Res. Microbiol. 144, 165–169 (1993).

38. Ruggiero, A. et al. A loose domain swapping organization confers a remarkable stability to the dimeric structure of the arginine binding protein from *Thermotoga maritima*. PLOS One 9, e96560 (2014).

39. Zhao, G., Xia, T., Fischer, R. & Jensen, R. Cyclohexadienyl dehydratase from *Pseudomonas aeruginosa*: Molecular cloning of the gene and characterization of the gene product. J. Biol. Chem. 267, 2487–2493 (1992).

40. Clifton, B. E. et al. Evolution of cyclohexadienyl dehydratase from an ancestral solute-binding protein. Nat. Chem. Biol. 14, 542–547 (2018).

41. Mahawaththa, M. C. et al. Small neutral Gd(III) tags for distance measurements in proteins by double electron-electron resonance experiments. Phys. Chem. Chem. Phys. 20, 23535–23545 (2018).

42. Goldfarb, D. Gd^3+^ spin labeling for distance measurements by pulse EPR spectroscopy. Phys. Chem. Chem. Phys. 16, 9685–9699 (2014).

43. Feintuch, A., Otting, G. & Goldfarb, D. Gd^3+^ Spin labeling for measuring distances in biomacromolecules: Why and how? in Electron Paramagnetic Resonance Investigations of Biological Systems by Using Spin Labels, Spin Probes, and Intrinsic Metal Ions, Part A (eds. Qin, P. Z. & Warncke, K. B. T.-M. in E.) 563, 415–457 (Academic Press, 2015).

44. Neylon, C. et al. Interaction of the *Escherichia coli* replication terminator protein (Tus) with DNA: a model derived from DNA-binding studies of mutant proteins by surface plasmon resonance. Biochemistry 39, 11989–11999 (2000).

45. Qi, R. & Otting, G. Mutant T4 DNA polymerase for easy cloning and mutagenesis. PLOS One 14, e0211065 (2019).

46. Mukai, T. et al. Highly reproductive *Escherichia coli* cells with no specific assignment to the UAG codon. Sci. Rep. 5, 9699 (2015).

47. Young, D. D. et al. An evolved aminoacyl-tRNA synthetase with atypical polysubstrate specificity. Biochemistry 50, 1894–1900 (2011).

48. Cabrita, L. D. et al. Enhancing the stability and solubility of TEV protease using in silico design. Protein Sci. 16, 2360–2367 (2007).

49. Abdelkader, E. H. et al. Protein conformation by EPR spectroscopy using gadolinium tags clicked to genetically encoded *p*-azido-L-phenylalanine. Chem. Commun. 51, 15898–15901 (2015).

50. Goldfarb, D. et al. HYSCORE and DEER with an upgraded 95 GHz pulse EPR spectrometer. J. Magn. Reson. 194, 8–15 (2008).

51. Mentink-Vigier, F. et al. Increasing sensitivity of pulse EPR experiments using echo train detection schemes. J. Magn. Reson. 236, 117–125 (2013).

52. Doll, A. et al. Gd(III)–Gd(III) distance measurements with chirp pump pulses. J. Magn. Reson. 259, 153–162 (2015).

53. Bahrenberg, T. et al. Improved sensitivity for W-band Gd(III)-Gd(III) and nitroxide-nitroxide DEER measurements with shaped pulses. J. Magn. Reson. 283, 1–13 (2017).

54. Bahrenberg, T., Yang, Y., Goldfarb, D. & Feintuch, A. rDEER: A modified DEER sequence for distance measurements using shaped pulses. Magnetochemistry 5, 20 (2019).

55. Jeschke, G. et al. DeerAnalysis2006—a comprehensive software package for analyzing pulsed ELDOR data. Appl. Magn. Reson. 30, 473–498 (2006).

56. Kabsch, W. XDS. Acta Crystallogr. D Biol. Crystallogr. 66, 125–132 (2010).

57. Evans, P. R. & Murshudov, G. N. How good are my data and what is the resolution? Acta Crystallogr. Sect. D Biol. Crystallogr. 69, 1204–1214 (2013).

58. McCoy, A. J. et al. Phaser crystallographic software. J. Appl. Cryst. 40, 658–674 (2007).

59. Emsley, P., Lohkamp, B., Scott, W. G. & Cowtan, K. Features and development of Coot. Acta Crystallogr. D Biol. Crystallogr. 66, 486–501 (2010).

60. Murshudov, G. N., Vagin, A. A. & Dodson, E. J. Refinement of macromolecular structures by the maximum-likelihood method. Acta Crystallographica Section D: Biological Crystallography 53, 240–255 (1997).

61. Afonine, P. V. et al. Towards automated crystallographic structure refinement with *phenix.refine*. Acta Crystallogr. Sect. D Biol. Crystallogr. 68, 352–367 (2012).

62. Bowers, K. J. et al. Scalable algorithms for molecular dynamics simulations on commodity clusters. in Proceedings of the ACM/IEEE SC Conference on Supercomputing (SC06), Tampa, Florida 43–43 (2007). doi:doi:10.1109/sc.2006.54

63. Roos, K. et al. OPLS3e: Extending force field coverage for drug-like small molecules. J. Chem. Theory Comput. 15, 1863–1874 (2019).

64. Bakan, A., Meireles, L. M. & Bahar, I. ProDy: Protein dynamics inferred from theory and experiments. Bioinformatics 27, 1575–1577 (2011).

65. Bennett, B. D. et al. Absolute metabolite concentrations and implied enzyme active site occupancy in *Escherichia coli*. Nat. Chem. Biol. 5, 593–599 (2009).

66. Campbell, E. et al. The role of protein dynamics in the evolution of new enzyme function. Nat. Chem. Biol. 12, 944–950 (2016).

67. Hong, N.-S. et al. The evolution of multiple active site configurations in a designed enzyme. Nat. Commun. 9, 3900 (2018).

68. Clifton, B. E. et al. Ancestral Protein Reconstruction and Circular Permutation for Improving the Stability and Dynamic Range of FRET Sensors. in Synthetic Protein Switches: Methods and Protocols (ed. Stein, V.) 71–87 (Springer New York, New York, 2017).

69. Whitfield, J. H. et al. Construction of a robust and sensitive arginine biosensor through ancestral protein reconstruction. Protein Sci. 24, 1412–1422 (2015).

70. Mitchell, J. A. et al. Rangefinder: A semisynthetic FRET sensor design algorithm. ACS Sensors 1, 1286–1290 (2016).

71. Fraser, N. J. et al. Evolution of protein quaternary structure in response to selective pressure for increased thermostability. J. Mol. Biol. 428, 2359–2371 (2016).

72. Marinelli, F. & Fiorin, G. Structural characterization of biomolecules through atomistic simulations guided by DEER measurements. Structure 27, 359-370.e12 (2019).

73. Stelzl, L. S., Erlenbach, N., Heinz, M., Prisner, T. F. & Hummer, G. Resolving the conformational dynamics of DNA with Ångstrom resolution by pulsed electron–electron double resonance and molecular dynamics. J. Am. Chem. Soc. 139, 11674–11677 (2017).

74. Puljung, M. C., DeBerg, H. A., Zagotta, W. N. & Stoll, S. Double electron - Electron resonance reveals cAMP-induced conformational change in HCN channels. Proc. Natl. Acad. Sci. U. S. A. 111, 9816–9821 (2014).

75. Joseph, B., Sikora, A. & Cafiso, D. S. Ligand induced conformational changes of a membrane transporter in E. coli cells observed with DEER/PELDOR. J. Am. Chem. Soc. 138, 1844–1847 (2016).

76. Joseph, B. et al. In situ observation of conformational dynamics and protein ligand–substrate interactions in outer-membrane proteins with DEER/PELDOR spectroscopy. Nat. Protoc. 14, 2344–2369 (2019).

