## Supplemental Data for "Altered conformational sampling along an evolutionary trajectory changes the catalytic activity of an enzyme"

* Correspondin­g authors.

**Table S1. Sites mutated for the incorporation of AzF residues.**

| Protein | Mutation sites^a^ |
| --- | --- |
| AncCDT-1 | Q68/K138; Q68/Q219; K138/S161 |
| AncCDT-3 | Q68/R138 |
| AncCDT-5 | Q68/R138 |
| *Pa*CDT^b^ | A68/A139 |

^a^ Amber stop codons were introduced to incorporate pairs of AzF residues. The positions in the amino acid sequence are indicated together with the original amino acid types.

^b^ The mutant A68/R138 was also prepared, but the tagging yields obtained were too low for DEER experiments.

**Table S2. Data collection and refinement statistics for the AncCDT-5 X-ray crystal structure.**

|  | **AncCDT-5 (HEPES)** |
| --- | --- |
| **PDB** | 6OKI |
| **Data collection** |  |
| Space group | P 6_5_ 2 2 |
| Cell dimensions |  |
| *a*, *b*, *c* (Å) | 70.74, 70.74, 174.28 |
| α, β, γ (°) | 90, 90, 120 |
| Resolution (Å) | 35.51–1.37 (1.42–1.37)* |
| *R*_merge_ | 0.08291 (4.382) |
| *R_pim_* | 0.020 (1.30) |
| *I*/σ*I* | 21.48 (0.57) |
| CC_1/2_ | 1.00 (0.338) |
| Completeness (%) | 99.67 (97.54) |
| Redundancy | 34.7 (28.3) |
| **Refinement** |  |
| Resolution (Å) | 35.51–1.37 (1.42–1.37) |
| No. reflections | 54894 (5266) |
| *R*_work_ / *R*_free_ | 0.1751 (0.3524) / 0.1954 (0.3580) |
| No. atoms | 2179 |
| Protein | 1884 |
| Ligand/ion | 17 |
| Water | 278 |
| *B*-factors | 33.05 |
| Protein | 31.58 |
| Ligand/ion | 38.62 |
| Water | 42.64 |
| R.m.s. deviations |  |
| Bond lengths (Å) | 0.006 |
| Bond angles (°) | 1.16 |

* Statistics for the highest-resolution shell are shown in parentheses.

**Table S3. Cα–Cα distances (tag positions) and calculated maxima of the Gd(III)–Gd(III) distance distributions for crystal structures and representative MD snapshots.**^a^

| **Name of Crystal Structure/MD Snapshot** | **Cα–Cα distance (nm)** | **Calculated Gd(III)–Gd(III) distance (nm)** |
| --- | --- | --- |
| **L-Arg AncCDT-1**  **(PDB 5T0W)** | **2.61** | **2.94** |
| MD_Anc1_closed_01 | 2.52 | 2.89 |
| MD_Anc1_closed_02 | 2.63 | 2.88 |
| MD_Anc1_closed_03 | 2.73 | 3.28 |
| MD_Anc1_closed_04 | 2.82 | 3.18 |
| MD_Anc1_closed_05 | 2.91 | 3.39 |
| **apo-AncCDT-1**  **(PDB 5TUJ)** | **3.36** | **4.06** |
| MD_Anc1_open_01 | 3.19 | 3.87 |
| MD_Anc1_open_02 | 3.40 | 4.90 |
| MD_Anc1_open_03 | 3.53 | 4.19 |
| MD_Anc1_open_04* | 3.63 | 4.10 |
| MD_Anc1_open_05 | 3.65 | 4.30 |
| MD_Anc1_wide_01 | 3.79 | 4.53 |
| MD_Anc1_wide_02 | 3.95 | 5.23 |
| MD_Anc1_wide_03 | 3.97 | 4.68 |
| MD_Anc1_wide_04 | 4.07 | 5.20 |
| MD_Anc1_wide_05 | 4.17 | 4.99 |
| **AncCDT-3/L188**  **(PDB 5JOS)** | **4.21** | **5.25** |
| MD_Anc3_L188_wide_01 | 4.04 | 5.03 |
| MD_Anc3_L188_wide_02 | 4.07 | 5.42 |
| MD_Anc3_L188_wide_03 | 4.26 | 5.54 |
| MD_Anc3_L188_wide_04 | 4.46 | 5.85 |
| MD_Anc3_L188_wide_05 | 4.67 | 6.24 |
| MD_Anc3_P188_wide_01 | 4.10 | 5.27 |
| MD_Anc3_P188_wide_02 | 4.27 | 5.57 |
| MD_Anc3_P188_wide_03 | 4.56 | 5.73 |
| MD_Anc3_P188_wide_04 | 4.68 | 6.22 |
| MD_Anc3_P188_wide_05 | 4.78 | 6.34 |

| **Name of Crystal Structure/MD Snapshot** | **Cα–Cα distance (nm)** | **Calculated Gd(III)–Gd(III) distance (nm)** |
| --- | --- | --- |
| **HEPES-AncCDT-5**  **(PDB 6OKI)** | **2.77** | **3.09** |
| MD_Anc5_closed_01 | 2.50 | 2.79 |
| MD_Anc5_closed_02 | 2.56 | 3.09 |
| MD_Anc5_closed_03 | 2.63 | 3.11 |
| MD_Anc5_closed_04 | 2.71 | 3.42 |
| MD_Anc5_closed_05 | 2.79 | 3.41 |
| MD_Anc5_open_01 | 3.01 | 3.40 |
| MD_Anc5_open_02 | 3.11 | 3.67 |
| MD_Anc5_open_03 | 3.23 | 4.00 |
| MD_Anc5_open_04 | 3.34 | 4.18 |
| MD_Anc5_open_05 | 3.44 | 4.17 |
| MD_Anc5_wide_01 | 3.56 | 4.54 |
| MD_Anc5_wide_02 | 3.68 | 4.52 |
| MD_Anc5_wide_03 | 3.93 | 5.12 |
| MD_Anc5_wide_04 | 4.06 | 5.16 |
| MD_Anc5_wide_05 | 4.21 | 5.44 |
| **HEPES-*Pa*CDT**  **(PDB 3KBR)** | **2.53** | **3.47** |
| **Acetate-*Pa*CDT**  **(PDB 5HPQ)** | **2.49** | **3.45** |
| MD_PaCDT_closed_01 | 2.58 | 3.49 |
| MD_PaCDT_closed_02 | 2.67 | 3.70 |
| MD_PaCDT_closed_03 | 2.76 | 3.84 |
| MD_PaCDT_closed_04 | 2.82 | 3.54 |
| MD_PaCDT_closed_05 | 2.96 | 3.85 |
| MD_PaCDT_open_01 | 3.15 | 3.99 |
| MD_PaCDT_open_02 | 3.33 | 4.06 |
| MD_PaCDT_open_03 | 3.59 | 4.40 |
| MD_PaCDT_open_04 | 3.61 | 4.46 |
| MDPaCDT_open_05 | 3.81 | 4.40 |
| **apo-*Tm*ArgBP**  **(PDB 4PRS)** | **4.00** |  |

^a^ Gd(III)–Gd(III) distances were calculated following modelling of the AzF-propargyl-DO3A-Gd(III) residue at each mutation site on structures obtained from crystallography (bold) or MD simulations. The models were built by identifying the most favourable values of the dihedral angles χ_1_ and χ_2_ of the tag (using the mutation tool of PyMOL (The PyMOL Molecular Graphics System, Version 2.0 Schrödinger, LCC.)) and setting χ_6_ to 180° as in the conformation depicted in Figure S2^1^. For the MD snapshots, structures with varying Cα–Cα distances (“MD_...._01-05”) were selected as representative structures from each conformational state (i.e. closed, open, wide-open). The data points representing these structures are shown as colored crosses in Figure 4.

*This snapshot could be grouped as being either open (based on the Cα–Cα distance) or wide-open (based on PCA analysis or radius of gyration), and so was included in both the open and wide-open groups.

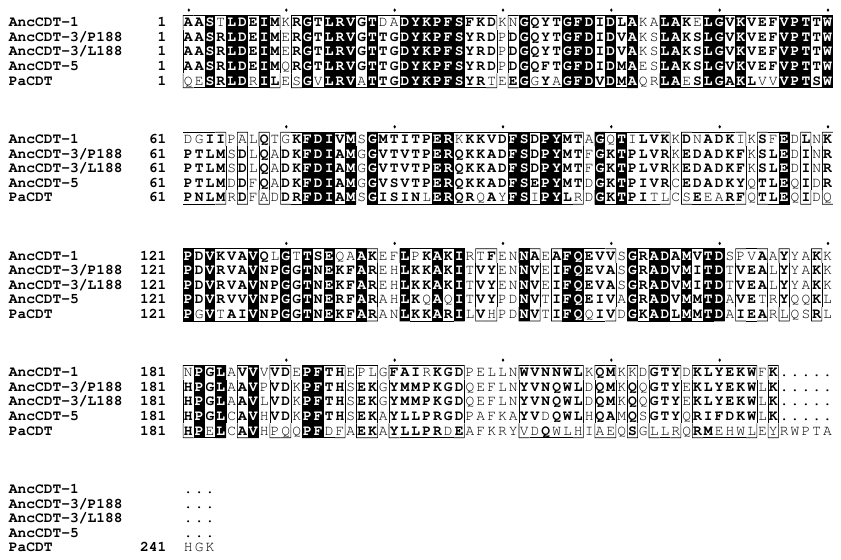

**Figure S1. Multiple sequence alignment.** Multiple sequence alignment of the proteins studied in this work. N-terminal residues that were introduced during cloning, including the His_6_-tag, have been omitted. The residue numbering corresponds to that used throughout this work. The sequences were aligned using Clustal Omega^2^. The figure was generated using ESPript 3^3^. Every 10^th^ residue position in the alignment is indicated by a black dot. Shading is based on the percentage of equivalent residues at that position (i.e. black = 100 % conserved).

**A**

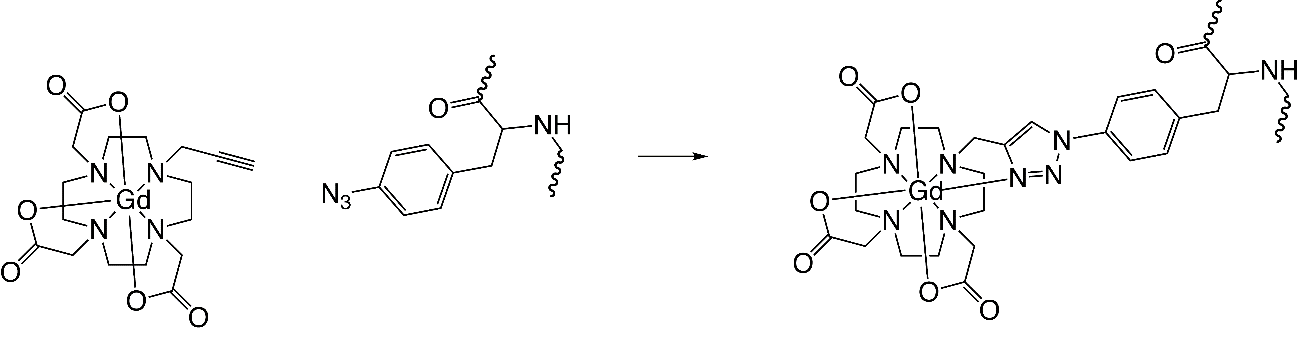

**B**

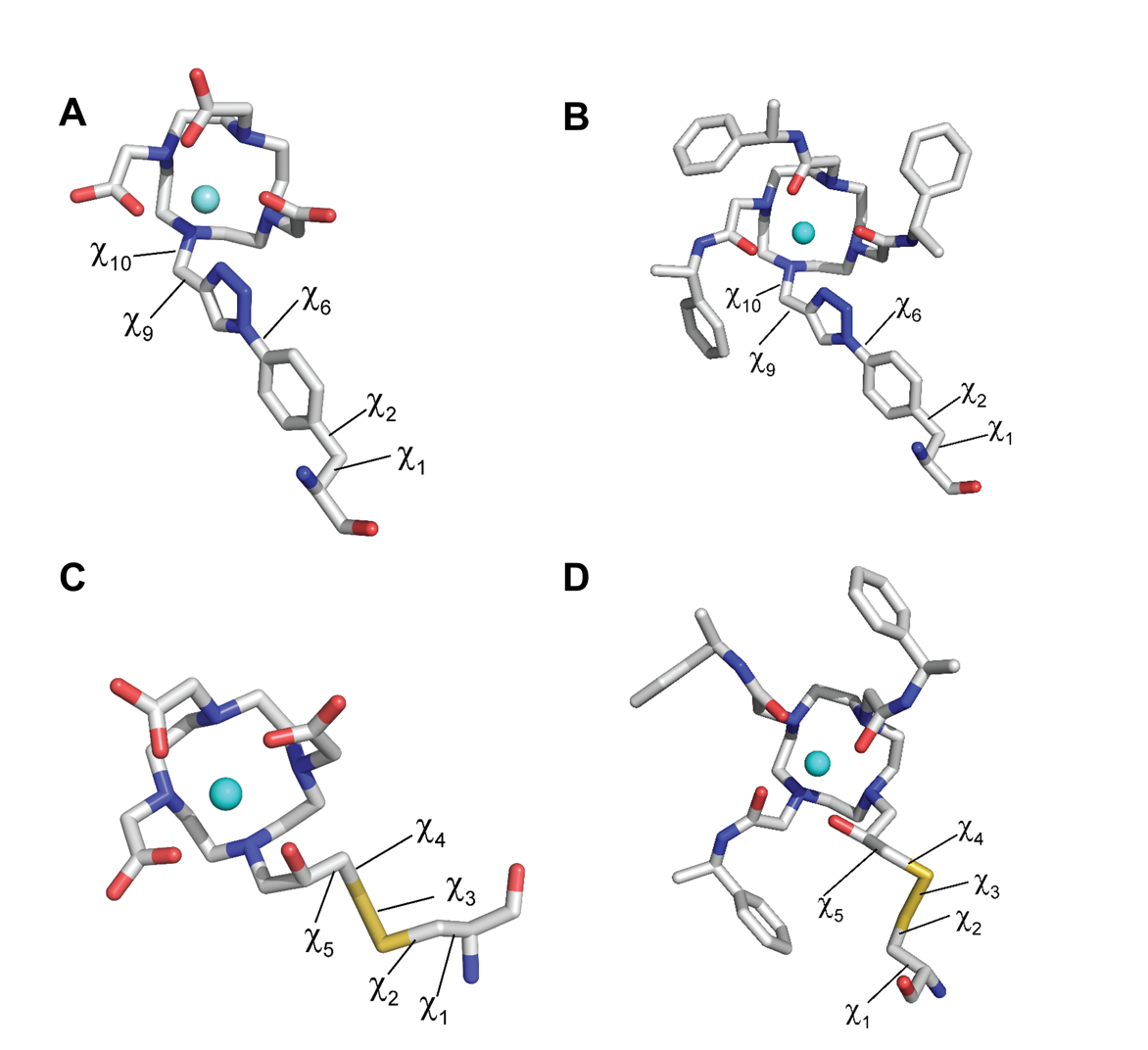

**Figure S2.** **The AzF-propargyl-DO3A-Gd(III) residue.** (A) Scheme of the Cu(I)-catalyzed cycloaddition reaction used to attach the propargyl-DO3A-Gd(III) tag to the AzF residue^4^. The reaction is biorthogonal and compatible with the presence of cysteine residues, as in AncCDT-5 and CDT. (B) Dihedral angles in the AzF-propargyl-DO3A-Gd(III) residue. χ_6_ = 0° in the conformation shown. χ_6_ = 180° was used for modelling in the present work, because using this value resulted in slightly better agreement between experimental and modelled Gd(III)–Gd(III) distances. The angles χ_9_ and χ_10_ were fixed to -140° and 70°, respectively, to allow coordination of the metal ion (cyan sphere) by the nearest nitrogen of the triazole ring.

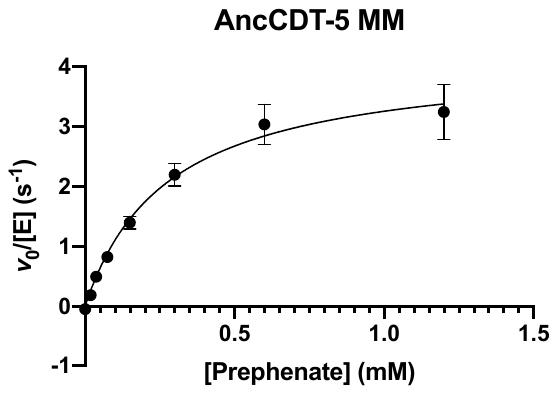

**Figure S3. Michaelis-Menten plot for AncCDT-5 prephenate dehydratase activity.** Results are mean ± s.e. of three technical replicates (n = 3).

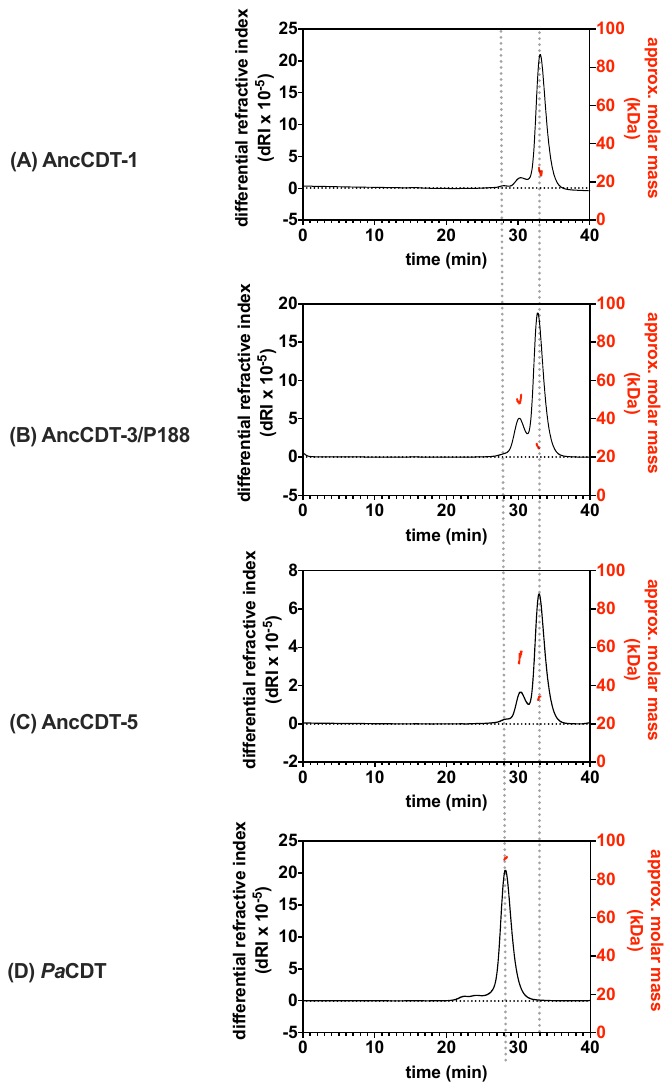

**Figure S4.** **SEC-MALS of proteins.** Results from analytical size-exclusion chromatography coupled with multiangle light scattering (SEC-MALS) for (A) AncCDT-1, (B) AncCDT-3/P188, (C) AncCDT-5 and (D) *Pa*CDT.

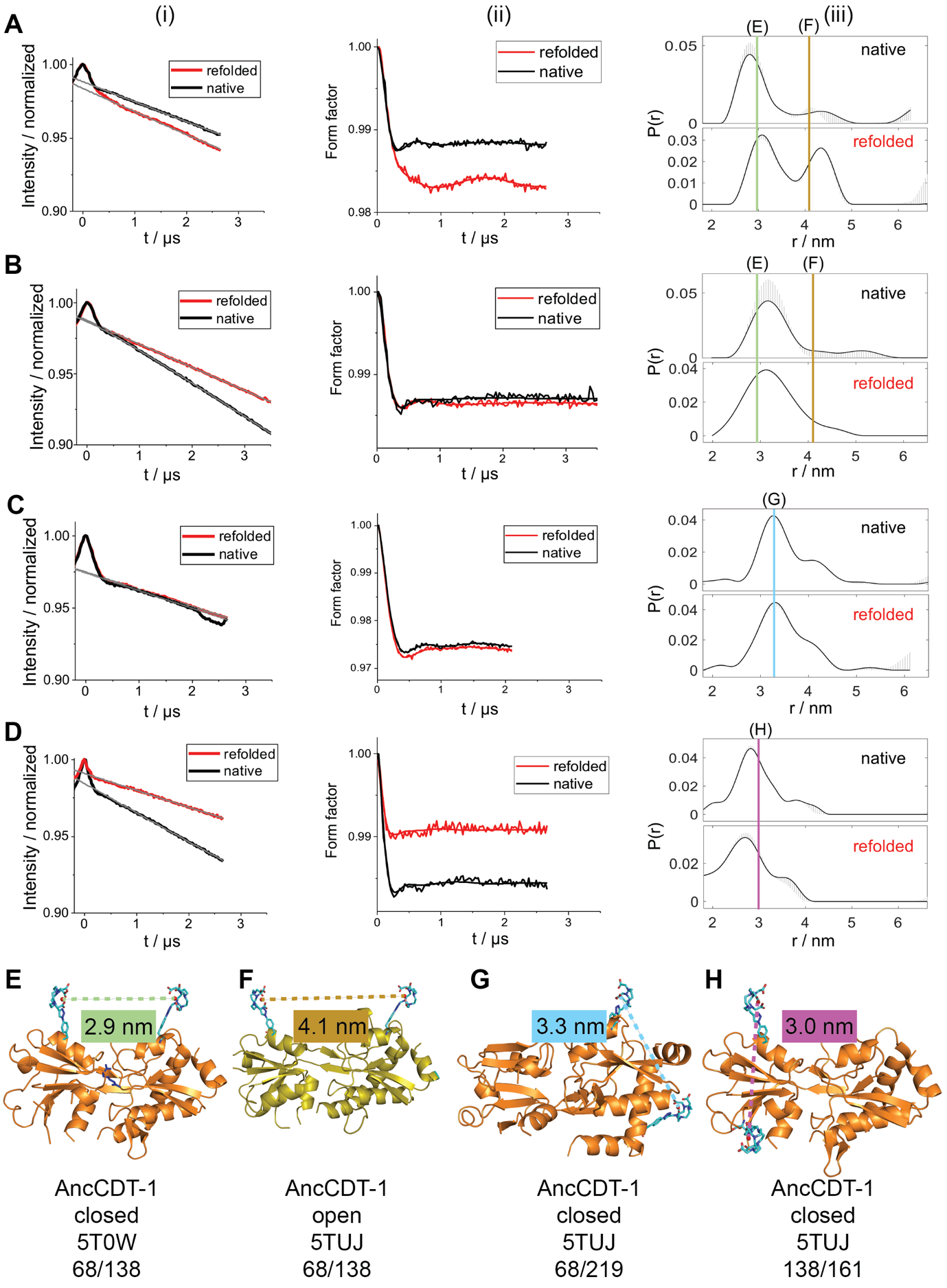

**Figure S5. Additional and primary DEER data for AncCDT-1.** Data from DEER measurements on natively purified (black) and refolded (red) AncCDT-1 samples (100 μM protein in D_2_O and 20 % w/v glycerol-d_8_) tagged at positions (A) 68 and 138 in the absence of exogenous L-Arg, (B) 68 and 138 in the presence of 1.5-fold molar excess L-Arg, (C) 68 and 219, or (D) 138 and 161. The left panel (i) shows the primary DEER data for the sample along with the fitted background (grey) function. The central panel (ii) shows the DEER form factors after background correction, with fits corresponding to the distance distributions shown in (iii). The right panel (iii) shows the corresponding distance distributions. The striped regions show alternative distributions obtained by varying the parameters of the background correction as calculated by the validation tool in the DeerAnalysis software package^5^. Vertical lines represent the predicted Gd(III)–Gd(III) distances when AzF-propargyl-DO3A-Gd(III) residues are modelled on crystal structures of (E) closed AncCDT-1 (PDB 5T0W) at positions 68 and 138, (F) open AncCDT-1 (PDB 5TUJ) at positions 68 and 138, (G) closed AncCDT-1 at positions 68 and 219, or (H) closed AncCDT-1 at positions 138 and 161.

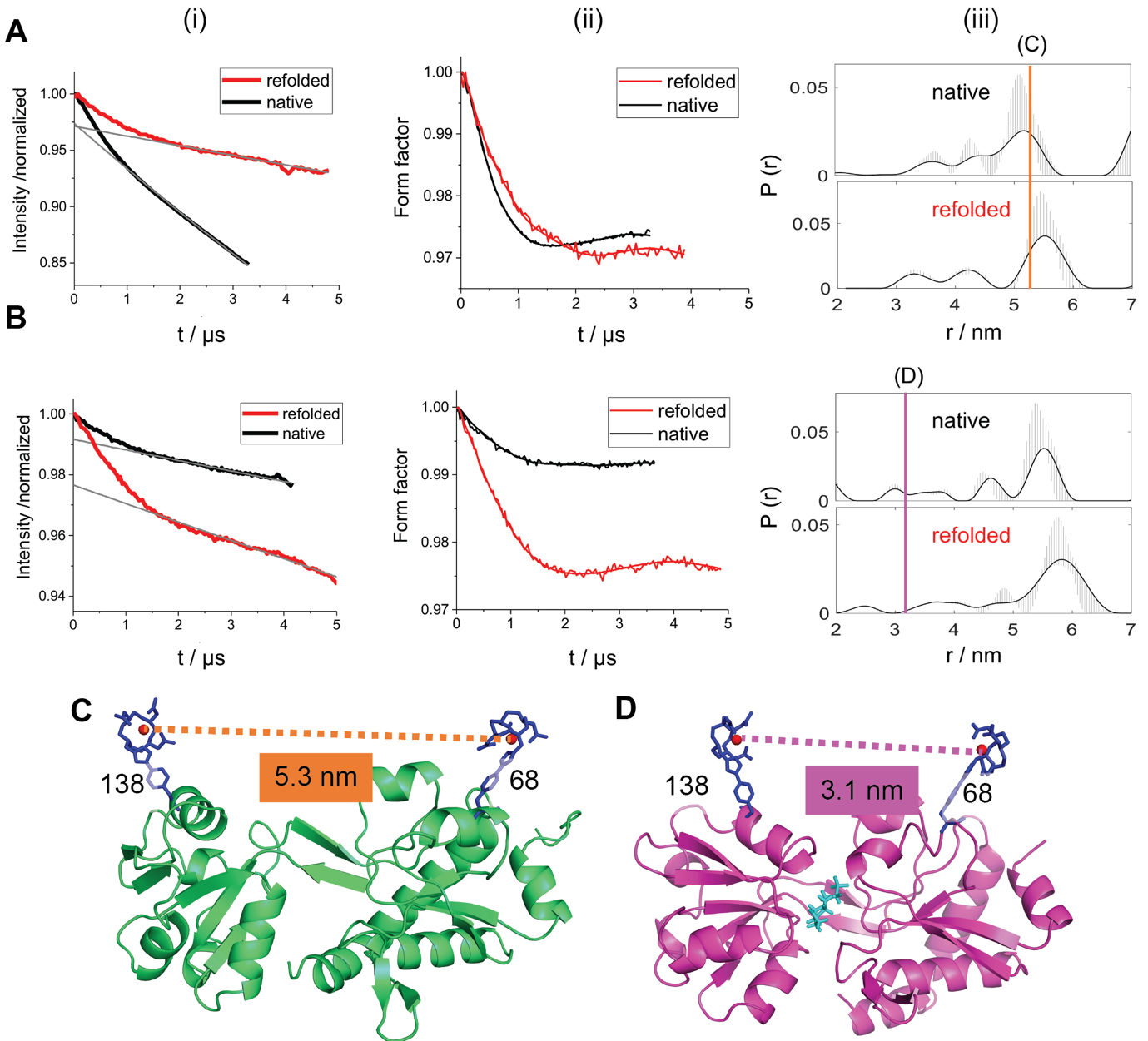

**Figure S6. Primary DEER data for AncCDT-3/P188 and AncCDT-5 samples.** Primary data from the DEER measurements (as shown in Figure 3) on natively purified (black) and refolded (red) of (A) AncCDT-3/P188 tagged at positions 68 and 138, and (B) AncCDT-5 tagged at positions 68 and 138. Samples were measured using 100 μM protein in D_2_O and 20 % w/v dglycerol-d_8_. The left panel (i) shows the primary DEER data for the sample and the fitted background (grey) function. The central panel (ii) shows the DEER form factors after background correction, with fits corresponding to the distance distributions shown in (iii). The right panel (iii) shows the corresponding distance distributions. The striped regions show alternative distributions obtained by varying the parameters of the background correction as calculated by the validation tool in DeerAnalysis^5^. Vertical lines represent the predicted Gd(III)–Gd(III) distances when AzF-propargyl-DO3A-Gd(III) residues are modelled onto crystal structures of (C) AncCDT-3/L188 (PDB 5JOS) at positions 68 and 138, or (D) AncCDT-5 (PDB 6OKI) at positions 68 and 138.

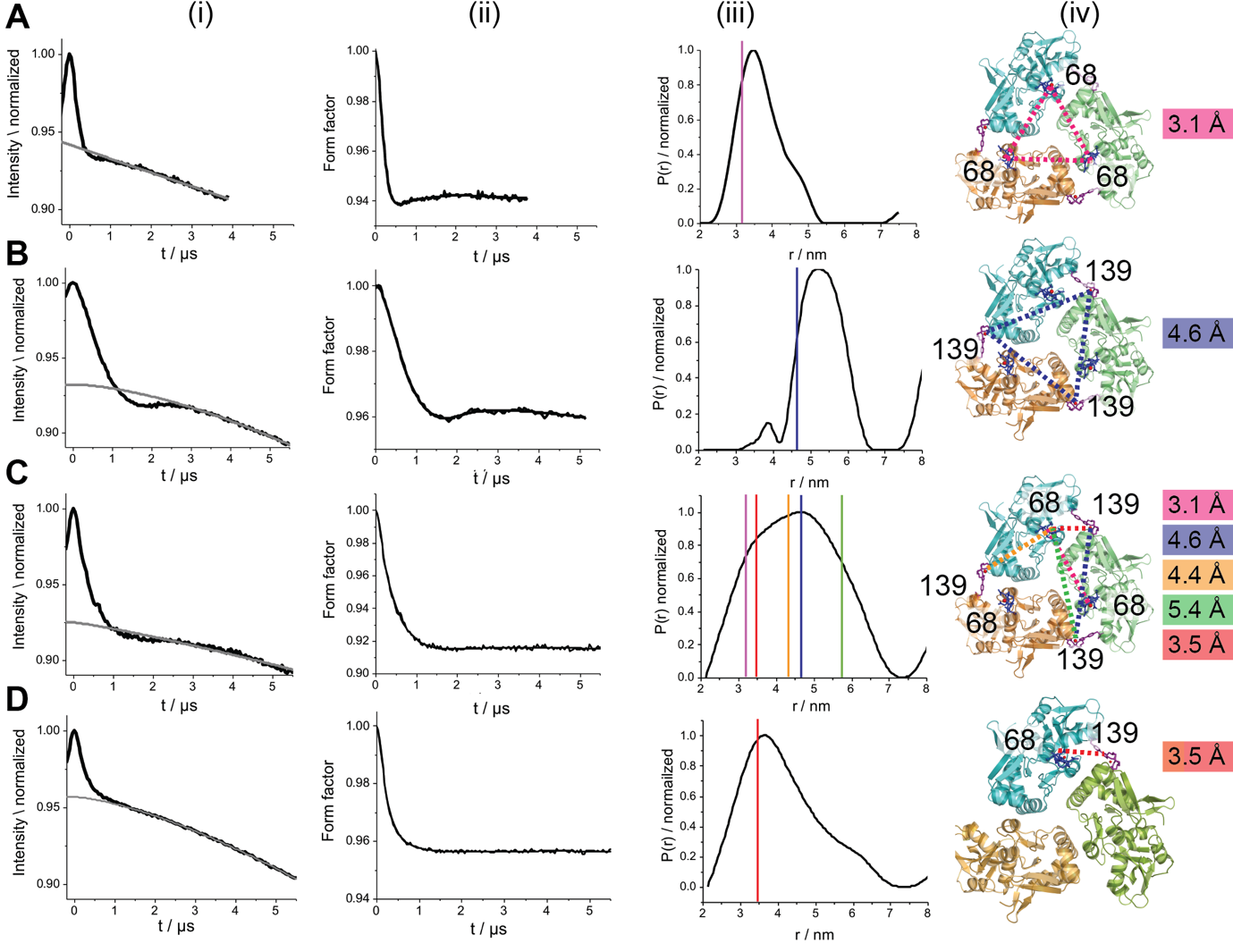

**Figure S7. Additional and primary DEER data for *Pa*CDT samples.** Data from the DEER measurements on natively purified samples of *Pa*CDT tagged at (A) position 68, (B) position 139, (C) position 68 and 139, or (D) position 68 and 139, but following a 10-fold dilution with unlabelled *Pa*CDT for selective observation of the intramolecular distance between the two domains (as shown in Figure 3). Samples were measured using 100 μM protein in D_2_O and 20 % w/v glycerol-d_8_. The left panel (i) shows the primary DEER data for the sample and the fitted background (grey) function. The central panel (ii) shows the DEER form factors after background correction, with fits corresponding to the distance distributions shown in (iii). The right panel (iii) shows the corresponding distance distributions. Vertical lines represent the predicted Gd(III)–Gd(III) distances when AzF-propargyl-DO3A-Gd(III) residues are modelled onto the crystal of structure *Pa*CDT (PDB 3KBR) at tagged positions, as shown in panel (iv).

**References**

1.	Mahawaththa, M. C. *et al.* Small neutral Gd(III) tags for distance measurements in proteins by double electron-electron resonance experiments. *Phys. Chem. Chem. Phys.* **20**, 23535–23545 (2018).

2.	Sievers, F. *et al.* Fast, scalable generation of high-quality protein multiple sequence alignments using Clustal Omega. *Mol. Syst. Biol.* **7**, 539 (2011).

3.	Robert, X. & Gouet, P. Deciphering key features in protein structures with the new ENDscript server. *Nucleic Acids Res.* **42**, W320–W324 (2014).

4.	Abdelkader, E. H. *et al.* Protein conformation by EPR spectroscopy using gadolinium tags clicked to genetically encoded *p*-azido-L-phenylalanine. *Chem. Commun.* **51**, 15898–15901 (2015).

5.	Jeschke, G. *et al.* DeerAnalysis2006—a comprehensive software package for analyzing pulsed ELDOR data. *Appl. Magn. Reson.* **30**, 473–498 (2006).
